# Hippocampal CA2 ripples recruit social replay and promote social memory

**DOI:** 10.1101/2020.03.03.975599

**Authors:** Oliva Azahara, Fernández-Ruiz Antonio, Leroy Felix, Siegelbaum A. Steven

## Abstract

The consolidation of spatial memory depends on the reactivation (‘replay’) of hippocampal place cells that were active during recent behavior. These reactivations are observed during sharp wave-ripples (SWRs), synchronous oscillatory events that occur during slow-wave sleep^1–9^ and whose disruption impairs spatial memory consolidation ^4,6,7,9^. Although the hippocampus encodes a wide range of non-spatial forms of declarative memory, it is not yet known whether SWRs are necessary for non-spatial memory. Moreover, although SWRs can arise from either the hippocampal CA3^8^ or CA2^10^ regions, the relative importance of these sources for memory consolidation is unknown. Here we examined the role of SWRs during the consolidation of social memory, the ability of an animal to recognize and remember a conspecific, focusing on CA2 because of its critical role in social memory^11,12,13^. We found that ensembles of CA2 pyramidal neurons that were active during social exploration of novel conspecifics were reactivated during SWRs. Importantly, disruption or enhancement of CA2 SWRs suppressed or prolonged social memory, respectively. Thus, SWR reactivation of hippocampal firing related to recent experience appears to be a general mechanism for binding spatial, temporal and sensory information into high-order memory representations.

---

We employed a social recognition task that assessed social memory formation in mice without interference from other social behaviors. In this task, a subject mouse was first habituated to a square arena with two empty wire cup cages in opposite corners. Immediately after habituation, we placed a novel stimulus mouse in each of the empty cups (mouse S_1_ and S_2_). In social learning trial 1 the subject mouse was allowed to explore the arena for 5 min. This was immediately followed by a second 5 min social learning trial with the positions of the same stimulus mice rotated 180° relative to trial 1 (Fig.1a, Extended Data Fig.1a,b). A one-hour sleep session preceded (pre-sleep) and followed (post-sleep) the two social learning trials. After the post-sleep period we exchanged one of the stimulus mice (chosen at random) for a third novel mouse (N) and performed a social memory recall trial.

**Figure 1:**
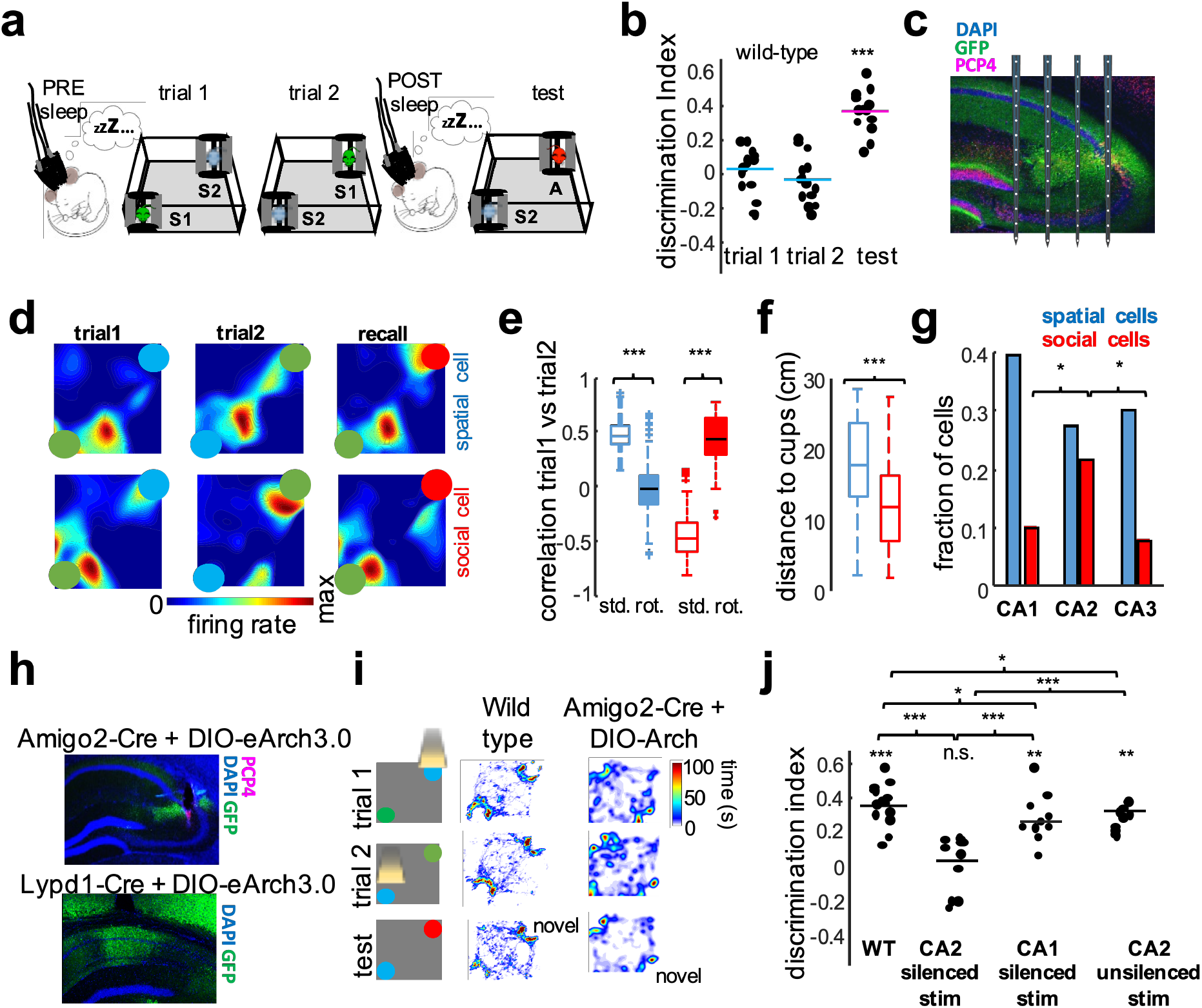
Encoding of conspecifics by CA2 pyramidal cells activity. **a**) Schema of the task. **b**) Task performance in wild-type mice (n = 13). Social memory quantified during recall trial by a discrimination index (see Methods). ***p < 0.0001, Wilcoxon sign-rank test. **c**) Histological verification of probe location. Note simultaneous electrodes in CA1, CA2 and CA3 regions. Blue, DAPI staining; green, GFP; red, PCP4 immunolabeling of CA2 pyramidal cells. **d**) Example firing maps of spatial and social CA2 place cells. **e**) Pearson’s correlation values for unrotated (std) and rotated (rot) place fields for spatial cells (n = 203 from 13 mice) and social cells (n=120 from 13 mice). Spatial cells had higher unrotated spatial correlations than rotated correlation (p < 10^−54^, sign-rank test); for social cells the reverse was true (p < 10^−16^). **f**) Social cells had place fields closer to the location of the stimulus mice than spatial cells (p < 10^−10^, rank-sum test). **g**) Proportion of spatial and social cells per region (CA1: 55 spatial /14 social; CA2: 131 /101; CA3: 17/5). Social cells were more abundant in CA2 than CA1 (p<0.05, Fisher’s test) or CA3 (p<0.05). **h**) eArch3.0-GFP expression (green) in CA2 pyramidal cells of an Amigo2-Cre mouse (top) and in CA1 cells of an Lypd1-Cre mouse (bottom). Blue, DAPI. **i**) Yellow light delivered to activate eArch3.0 during exploration of either S_1_ or S_2_ during trials 1 and 2. **j**) Silencing of CA2 impaired memory of stimulus mouse around which light was applied (n =10 mice; p<0.0001, sign-rank test) but not memory of stimulus mouse where light was not applied (distractor); CA1 silencing had no effect (n = 10; p > 0.05).

Animals interacted for equal times with S_1_ and S_2_ in the two learning trials and showed a significant preference (based on a social discrimination index, see Methods) for the novel animal introduced in the test trial (Fig.1b; p < 0.001 sign-rank test), with no significant differences in other behavioral parameters (Extended Data Fig.1c-d). The preference for the novel mouse demonstrated the successful encoding and consolidation of social memory of the stimulus mouse. To explore the activity of different hippocampal regions during this task, we implanted high-density silicon probes in the dorsal hippocampus (n=13; Fig.1c, Extended Data 2 and Table 1) and analyzed the spatial firing patterns of single pyramidal cells during the task (140 CA1 neurons from 8 mice; 473 CA2 neurons from 13 mice; 64 CA3 neurons from 5 mice).

**Table 1:** Description of animal cohorts included in the study

| **Cohort** | **Virus** | **Implant** | **Protocol** |
| --- | --- | --- | --- |
| 1 (amigo2-cre +)<br>n=7 | AAV5-DIO-Chr2-EYFP | 5X12-channel probes or<br>4x16 probes and 2 fibers | No stim., SWR disruption,<br>random disruption |
| 2 (amigo2-cre -) n=6 | AAV5-DIO-Chr2-EYFP | 5X12-channel probes or<br>4x16 probes and 2 fibers | No stim., SWR disruption |
| 3 (amigo2-cre +)<br>n=3 | AAV5-DIO-eArch3.0-EYFP | 5X12-channel probes<br>and 2 fibers | No stim., 30 s silencing (every<br>two minutes) |
| 4 (amigo2-cre +)<br>n=11 | AAV5-DIO-eArch3.0-EYFP | Optic fibers | No stim., 30 s silencing,<br>conspecific silencing, distractor<br>silencing |
| 5 (amigo2-cre -)<br>n=10 | AAV5-DIO-eArch3.0-EYFP | Optic fibers | No stim., 30 s silencing,<br>conspecific silencing |
| 6 (Lypd1-cre +) n=8 | AAV5-DIO-eArch3.0-EYFP | Optic fibers | No stim., 30 s silencing,<br>conspecific silencing |
| 7 (Grik4-cre +)<br>n=6 | AAV5-DIO-Chr2-EYFP | 5X12-channel probes or<br>4x16 probes and 2 fibers | No stim., SWR generation |
| 8 (Grik4-cre -)<br>n=6 | AAV5-DIO-Chr2-EYFP | 5X12-channel probes or<br>4x16 probes and 2 fibers | No stim., SWR generation |

**Table 2:**
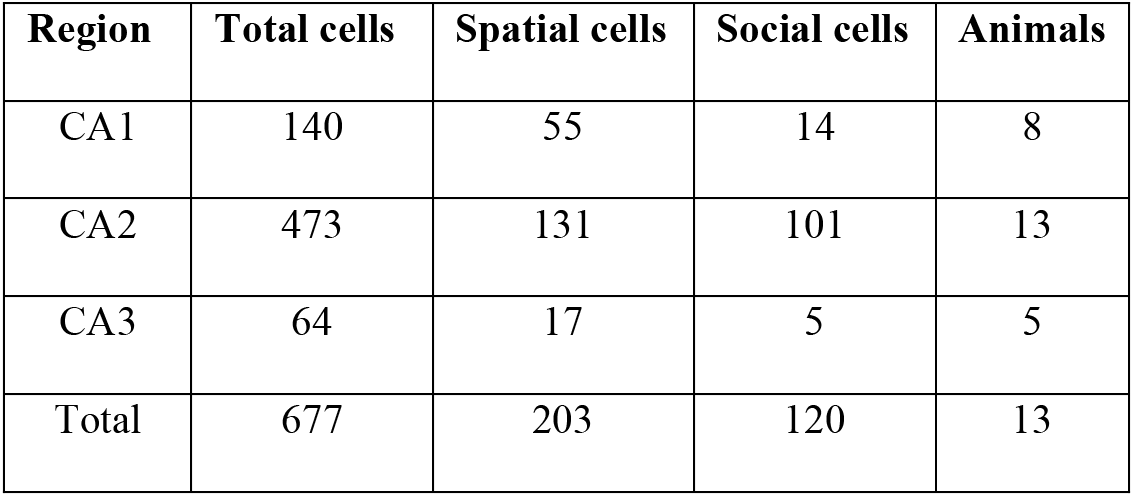
Number of cells recorded

We found two classes of neuronal responses. Some cells, which we termed “spatial cells”, displayed stable place fields relative to the arena across the three trials (Fig.1d, upper). In contrast, other cells, termed “social cells”, showed place fields that remapped between trial 1 and trial 2 in a manner that maintained firing field position relative to the location of one of the two stimulus mice (Fig.1d, bottom). We quantified this phenomenon by comparing the spatial correlation of firing maps between the two learning trials with the correlation obtained by rotating the map in trial 2 by 180° to match the position of the stimulus mice in trial 1 (Extended Data Fig.3).

Unsupervised clustering of a scatterplot of the unrotated versus rotated correlation values for all CA2 neurons revealed several clusters. One subpopulation of neurons (28% of the cells) had maximal unrotated and minimal rotated correlation values, which we defined as the spatial cells. A second subpopulation (21% of the cells) had maximal rotated and minimal unrotated correlation values, which we defined as the social cells (Fig.1e and Extended Data Fig.3c-d). Place fields of social cells lay closer to the cups containing stimulus mice (Fig.1f) but these cells otherwise displayed similar properties to spatial cells (Extended Data Fig.3e-i) Social cells were significantly more abundant in CA2 than in CA1 or CA3 (Fig.1g), consistent with the selective role of dorsal CA2 in social memory^11,12,13^.

To explore the contribution of CA1 and CA2 cells to social behavior in this task, we expressed in a Cre-dependent manner the inhibitory opsin archaerhodopsin (AAV-DIO-eArch3.0) in CA2 or in CA1 using the CA2-selective Amigo2-Cre mouse line or the CA1-selective Lypd1-Cre line, respectively. The animals were then implanted with bilateral optic fibers over dorsal CA2 or dorsal CA1 (Fig. 1h). We silenced CA2 or CA1 pyramidal cells by applying yellow light during the interaction times with either S_1_ or S_2_ (chosen at random for each experiment) during the two learning trials (Fig.1i). We found that silencing of CA2 but not CA1 disrupted social memory of the stimulus mouse around which light pulses were applied but did not affect memory of the other stimulus mouse (Fig.1j). These results extend previous reports on the importance of CA2 for social memory formation^11,12,13^ by showing the specificity for CA2 activity in learning the social identity of a given animal.

Next, we explored the neural substrate of social memory consolidation in the social recognition task. We observed a significant increase in the rate of SWRs during sleep after the social experience (post-sleep) compared to sleep before the social experience (pre-sleep; Fig.2a), similar to the increase in SWRs in rodents during sleep following spatial exploration^14^ and in humans engaged in a visual memory task^15,16,17^. We thus hypothesized that SWRs are involved in social memory consolidation. To test this idea, we performed closed-loop optogenetic disruption of SWRs during the consolidation stage (post-sleep) of the task in Amigo2-Cre mice expressing ChR2 in CA2 (Fig.2b). Strong photostimulation of these neurons with a brief intense light pulse upon SWR detection^9^ resulted in the premature termination of SWRs, with a period of post-excitation inhibition of CA2 firing (Fig.2c, Extended Data Fig.4a-b). In control sessions, we delivered the same light pulse after a random delay upon SWR detection (Fig.2c).

**Figure 2:**
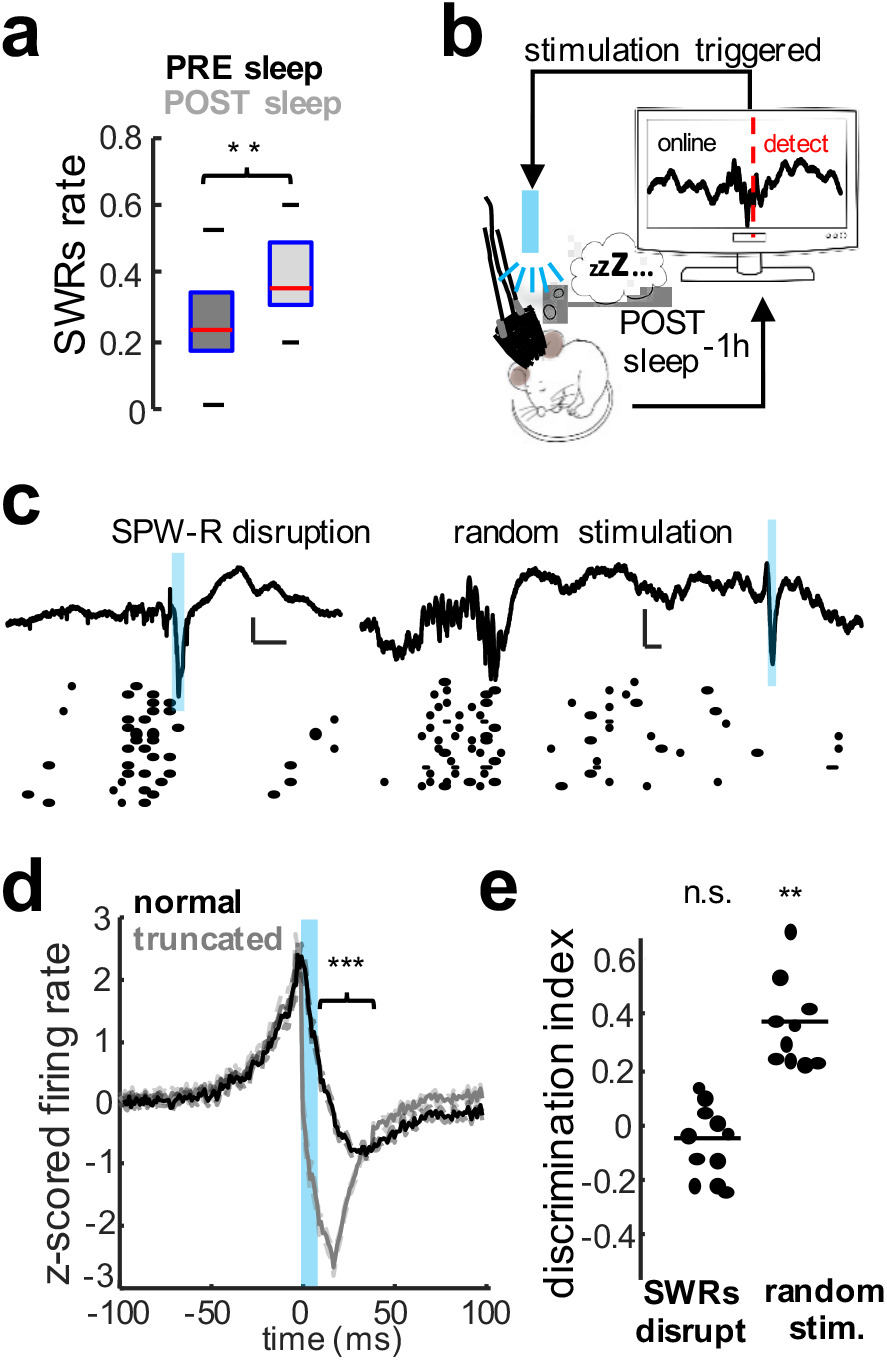
Effect of sharp-wave ripple disruption on social memory consolidation. **a**) SWR rate in CA2 increased during post-sleep compared to pre-sleep sessions (n = 17 mice; p = 0.007, rank-sum test). **b**) Schematic of SWRs disruption during sleep consolidation periods. **c**) ChR2 strongly activated with10-ms high-intensity blue light pulses upon real-time detection of ripples in CA2. Examples of closed-loop ripple truncation and random delay stimulation. Traces show LFPs from the CA2 pyramidal layer (top) and raster plot firing of individual CA2 units (bottom). **d**) Averaged firing rate response of all pyramidal cells to spontaneous and truncated SWRs. Note strong suppression of firing after SWR truncation compared to spontaneous SWR (p < 10^−4^, rank-sum test). **e**) Social memory recall was impaired following CA2 ripple disruption (p > 0.05, n = 7 mice) but preserved following random stimulation (p = 0.002, n = 7).

We found that SWR disruption during post-sleep impaired social memory recall in Amigo2-Cre animals expressing ChR2 whereas random stimulation had no effect (Fig.2d; n = 7). As an alternative approach, we directly silenced CA2 neurons with photoactivation of eArch3.0 for 30 s once every two minutes during the post-sleep period (Extended Data Fig.4c-d). This manipulation reduced the number of SWRs and significantly impaired social memory recall (Extended Data Fig.4e-f). Together, these results demonstrate that CA2 SWRs are necessary for social memory consolidation. Of note, in contrast to the inhibitory effects of relatively brief periods of CA2 silencing we observe, longer periods of CA2 silencing can increase SWRs^18^, likely as a result of disinhibition of CA3^19^.

SWRs during sleep following spatial learning have been shown to reactivate place cells that fired during spatial exploration^1,2,5,20,21^. To explore whether social firing was reactivated during SWRs, we adopted the explained variance (EV) approach previously used to assess reactivation of CA1 neurons following spatial exploration^21,22^. The EV measures the extent to which the variance in firing rate correlations between all pairs of neurons during SWRs in the post-sleep session could be explained by the correlations in firing of the same pairs of neurons during the preceding social exploration sessions, taking into account preexisting correlations observed in pre-sleep (Fig. 3a, Extended Data Fig. 5a; Methods)^21,22^. As a measure of chance levels of EV, we calculated the reverse explained variance (REV), the extent to which the variance in correlations during SWRs in pre-sleep could be explained by the variance during social exploration, taking into account the variance in correlations during the post-sleep period.

**Figure 3:**
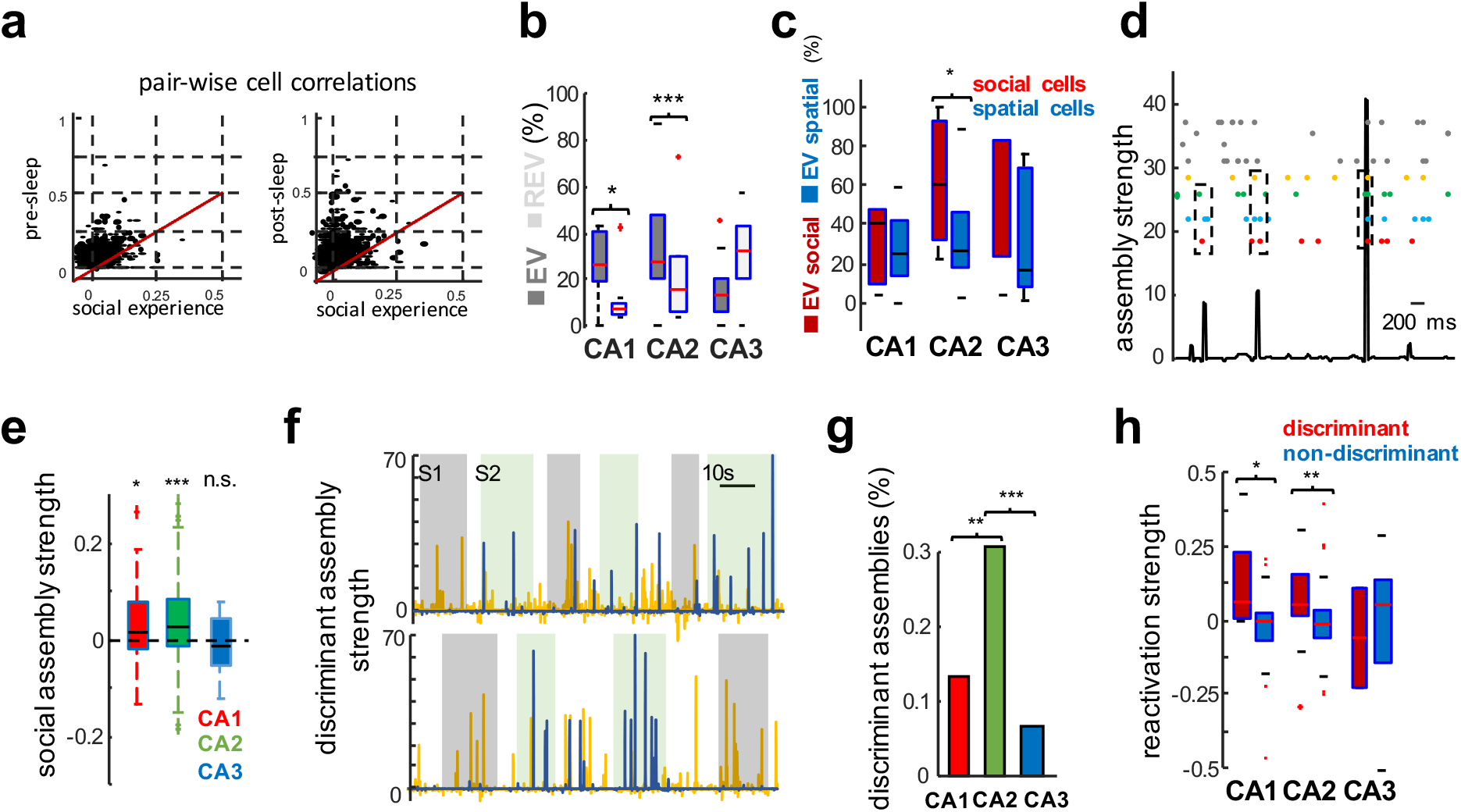
Reactivation during sleep of cell ensembles active during prior social learning. **a**) Pearson’s correlation coefficients for all pairs of neurons measured during pre-sleep (left) or post-sleep (right) versus correlations measured during social learning trials. **b**) EV (based on r values) was significantly higher than REV (chance) for CA2 (p = 0.0028, Wilcoxon sign-rank test, n = 13 mice) and CA1 (p = 0.03, n = 13 mice) cells but not CA3 (p > 0.05, n = 7 mice). **c**) EV for social versus spatial cells in each area (p < 0.05 for CA2 only, sign-rank test). **d**) Example of assembly detection. Each line of the raster plot shows the firing of one unit; units that are part of the example assembly are colored. Bottom trace shows activation strength for example assembly. **e**) Mean normalized social assembly strength (difference between assembly strength inside and outside social interaction zones divided by sum of assembly strengths) was significantly greater than 0 in CA1 (p < 10^−6^, sign-rank test) and CA2 (p < 10^−19^) but not CA3 (p > 0.05). **f**) Activation of discriminant assemblies. Recording showing one CA2 assembly (yellow line) more active during interactions with mouse S1 (grey squares) and a second (blue line) more active during interactions with mouse S2 (green squares). **g**) Fraction of discriminant assemblies in each region (p < 0.0034 and 10^−4^ for CA2 versus CA1 and CA3, respectively, Fisher’s test). **h**) Assembly reactivation strength (difference between assembly strength during post-sleep and pre-sleep) was higher for discriminant compared to non-discriminant assemblies in CA1 (p=0.043, rank-sum test) and CA2 (p = 0.0032) but not CA3 (p = 0.6837).

Based on this method, we found significant levels of reactivation in CA1 and CA2, but not CA3, during post-sleep SWRs (Fig. 3b). Moreover, optogenetic disruption of SWRs, but not random stimulation, abolished this reactivation (Extended Data Fig.5b). We also observed significant levels of reactivation when we calculated EV during firing throughout the entire slow wave sleep period, not just during SWRs. However, there was no significant reactivation during REM sleep epochs (Extended Data Fig.5c-d), in line with previous reports^21,22^. Of further interest, social CA2 cells had higher EV values than spatial cells in either CA2 or other areas (Fig.3c and Extended Data Fig.5), indicating that reactivations in CA2 preferentially represented social information.

We hypothesized that groups of cells that were co-active during a social exploration episode may form functional ensembles that are reactivated during SWRs, in a similar manner to reactivation of place cell ensembles representing experienced trajectories^1,2,3,5,7,9^. To test this idea, we identified a set of cell assemblies of co-active neurons during social exploration, using principal component analysis to first estimate the number of assemblies followed by independent component analysis to assign weights for each neuron’s contribution to a given assembly (component). Based on these weights and the z-scored firing rates of the neurons, we determined the strength of activation of each assembly as a function of time during a given session, as previously described^20,23^ (Fig.3d). When examined during the social learning trials, CA2 and CA1, but not CA3, cell assemblies were more strongly activated during periods of social interaction compared to other times during the trials (Fig.3e, Extended Data Fig.6b). Of particular interest, some assemblies, which we termed “discriminant assemblies”, were activated significantly more strongly during interactions with one stimulus mouse compared to the other (Fig.3f and Extended Data Fig.7b). Such discriminant assemblies were more abundant in CA2 than in CA1 or CA3 (Fig.3g).

Next, we explored whether cell assemblies identified during the social learning trials were reactivated during post-sleep SWRs and/or during social interactions during the recall trial. To distinguish activity of preexisting assemblies from activity resulting from social learning, we compared assembly activity in the post-sleep to pre-sleep sessions. We found significant reactivation of assemblies in all three hippocampal regions. Moreover, the strength of reactivation of discriminant assemblies during post-sleep and recall trials was stronger than that of non-discriminant assemblies (Fig.3h; Extended Data Fig.7c). Finally, peri-ripple assembly activation showed a higher activity of discriminant assemblies during SWRs in the sleep session after social learning than before (Extended Data Fig.8). These results suggest that hippocampal cell ensembles, particularly from the CA2 region, encode and help consolidate the memory of conspecifics.

Optogenetic stimulation with appropriately shaped light-pulses of CA1 pyramidal cells expressing ChR2 has been found to generate artificial ripple oscillations with features similar to spontaneous ripples^9,24,25^. Furthermore, optogenetic prolongation of CA1 SWRs during a spatial working memory task can improve memory performance^9^. Thus, to explore further whether CA2 reactivation during SWRs was important for social memory consolidation, we asked whether optogenetic generation of SWRs in CA2 in the post-sleep period after social learning could enhance social memory recall. We injected Cre-dependent AAV to express ChR2 in CA2 neurons of Amigo2-Cre mice, with a control group of Cre-littermates injected with the same virus and receiving the same light pulses.

Under the conditions of our social memory task, a subject mouse exhibited significant social memory after a 1-h but not a 24-h delay interval (Fig.4a). We applied weak (1-1.5 mW), 60-70-ms shaped light pulses to trigger ripples with features similar to spontaneous events (Fig.4c^9,24,25^), producing a ~2-fold increase in the number of ripples in the first 2 h of the 24-h delay period relative to controls (Extended Data Fig.9b). Both the firing rate and probability that a neuron fires at least one spike during SWRs (probability of participation) was similar for spontaneous compared to artificial ripples for CA1, CA2, and CA3 (Fig.4d, Extended Data Fig.9c-d), in agreement with prior results in CA1^9,24,25^. Importantly, mice that were stimulated to generate additional CA2 ripples during the post-sleep period showed a significant improvement in social memory when tested 24 h after social exposure (n=7; Fig.4e). In comparison, there was no effect of the light pulses on social memory in the control group of mice (n=6; Extended Data Fig.9e).

**Figure 4:**
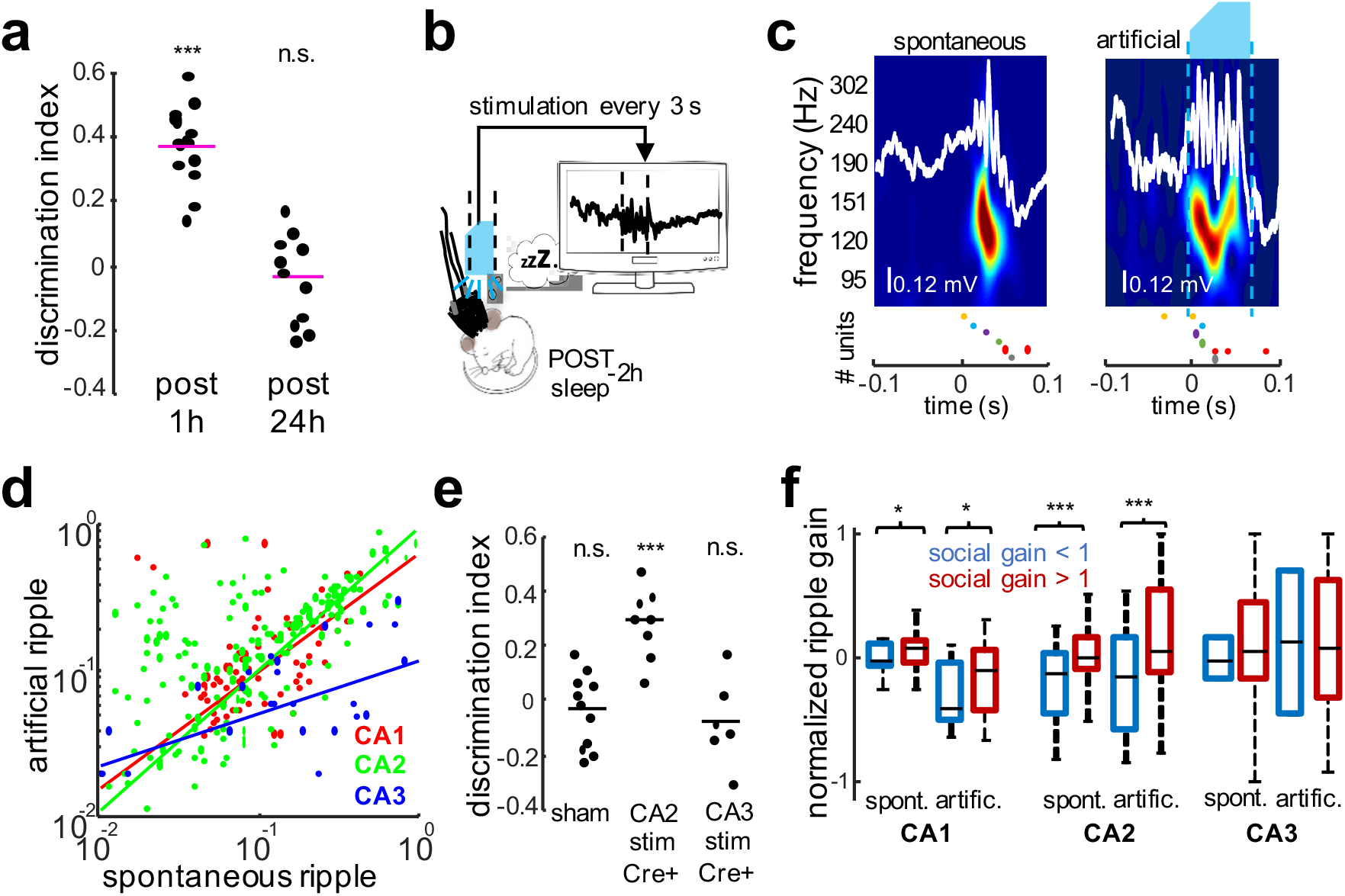
Effect of optogenetic generation of ripples on social memory recall. **a**) Social memory recall was normally absent 24 h after social learning (n = 7 mice; p > 0.05, sign-rank test). **b**) Weak ChR2 activation in CA2 neurons of Amigo2-Cre mice with low intensity blue light pulses induced artificial ripples in CA2 during the first 2-h of the post-sleep session. **c**) Example of spontaneous and artificial CA2 ripples and associated spiking from the same animal. White lines show CA2 LFPs, color maps show wavelet spectrogram, blue shape indicates light stimulus. The firing of each unit is color-coded in the two raster plots. **d**) Correlation of probability of participation in spontaneous and artificial CA2 ripples for CA1 (n = 67; r= 0.56, p < 10^−3^), CA2 (n= 147; r = 0.66, p < 10^−25^) and CA3 (n = 40; r = 0.26, p > 0.05) pyramidal cells. **e**) Social memory with 24 h delay in mice receiving blue light pulses for control group (Cre-littermates; p > 0.05, sign-rank test, n = 6 mice), Amigo2-Cre mice expressing ChR2 in CA2 (p < 0.00023, n = 6 mice), and Grik4-Cre mice expressing ChR2 in CA3 (p > 0.05, n = 6 mice). **f**) Ripple reactivation gain (firing rate increase over baseline during post-sleep SWRs – firing rate increase during pre-sleep SWRs)/(sum of two rates) for both spontaneous and artificial SWRs was greater for cells with positive social gain than those with negative social gain in CA1 (p<0.05, n=67) and CA2 (p<0.003, n=147) but not CA3 (p>0.05, n=40).

We next asked whether the socially-related firing of cells during the social learning trials influenced their reactivation in spontaneous and artificial ripples. We classified the social firing of each cell by its “social gain” in the social learning trials: (firing rate during social interactions - firing rate outside of social interactions)/(sum of the rates). Indeed, we found that cells with a positive social gain had a greater ripple reactivation gain, the extent to which firing rate during post-sleep ripples was increased relative to pre-sleep ripples (Fig.4f). Cells with positive social gain also displayed a higher probability of participation in SWRs during spontaneous and artificial events (Extended Data Fig.9f). Previously, it has been shown that synaptic plasticity can be induced by pairing stimulation of CA3 inputs onto CA1 with SWRs events^26^ and by pairing local stimulation with naturally occurring SWRs^27^. Our data suggest that artificial generation of SWRs via optogenetic stimulation it is able to potentiate previously active synapses. In addition, it was recently suggested that SWRs are privileged windows for learning-related changes in excitability^28^, and that a novel experience or learning preferentially recruits new cells, which in turn, due to intrinsic or synaptic plasticity properties, are more highly reactivated during SWRs after learning^5,9,28^. Our results, support this idea by finding that social cells were highly recruited during SWRs after the social episode (Extended Data Fig.6).

Finally, we examined whether the effect of SWRs to enhance social memory was specific to SWRs generated by CA2, or whether SWRs triggered in neighboring CA3 were also sufficient. We thus expressed ChR2 in CA3 pyramidal neurons, using Cre-dependent AAV injections in CA3 of Grik4-Cre mice, which express Cre in CA3 pyramidal cells^29^. Although photostimulation of CA3 was able to successfully generate SWRs (Extended Data Fig.10c-d), this manipulation failed to improve social memory recall after 24 h (Fig.4e; Fig.10a-b; n = 6). Of interest, CA3-triggered SWRs recruited the firing of pyramidal neurons in CA3 and CA1 but not in CA2 (Extended Data Fig.10e-g), suggesting that CA2 firing activity during SWRs is critical for social memory consolidation. This finding is consistent with results showing that dorsal CA2 but not dorsal CA1 or CA3 is important for social memory^11,12,13,30^, and that CA3 excitatory inputs to CA2 recruit little net excitation due to strong feedforward inhibition^31^.

Our results demonstrate that the cellular substrate for social memory consolidation in the dorsal hippocampus relies on SWRs generated in dorsal CA2 but not CA3. We found signatures of social identity encoding in the dorsal CA2 region at both the single cell (Fig.1) and cell ensemble levels (Fig.3), consistent with findings that CA2 is necessary for social memory^11,12,13^ and that CA2 firing responds to social novelty^13,30^. In addition, we also observed that CA1 pyramidal neuron firing contains significant social information (Fig.1). Although dorsal CA1 does not appear to be required for social memory^11^, the social information in dorsal CA2 may enable certain CA1 neurons to act as social place cells, firing in response to the specific location of a conspecific^32,33^. Previous reports showed that hippocampal place cells fired in response to a continuously viewed external stimulus^34,35^. In these studies, a subgroup of cells showed a stable response, others rotated with the non-spatial cue and others showed a less specific and unstable response. Our results are in line with these studies and add the fact that 1) the receptive fields of “place-cells” can respond to higher order external variables, such as the identity of particular conspecifics, 2) “place-cells” that responded to conspecifics participated with higher probability during post-experience SWRs. In addition, our finding that hippocampal neurons may respond to particular conspecifics is consistent with a previous report that individual neurons in the medial temporal lobe of humans respond to the identity of given individuals^36^. Importantly, our results provide the first causal evidence that SWRs are necessary for non-spatial memory consolidation. Furthermore, our findings suggest that SWRs originating from different hippocampal regions may have different functional roles. Thus, whereas acute silencing of CA3 reduces SWRs and the firing responses of CA1 cells during spatial memory tasks^37^, consolidation of social memory requires SWRs arising selectively in CA2. The idea that CA2 SWRs may participate in social memory is also consistent with the finding that the social neuropeptides oxytocin and vasopressin can heighten CA2 intrinsic^38^ and extrinsic^39^ excitation. On the other hand, it has been recently shown that a functional circuit from dCA2 to vCA1 is involved in social memory^12^ and that the rate of SWRs in vCA1 is increased by the presence of a conspecific^40^. These findings together with our current results suggest that SWRs originated from CA2 might propagate to vCA1 in order to consolidate the social memory engram. This idea is further supported by the findings that vCA1 also store conspecific information^41^. Furthermore, our finding that cells and assemblies that encoded social identity during experience were preferentially reactivated during post-experience SWRs (Fig.3) suggests a general role for SWRs in orchestrating the reactivation of recently experienced social information by patterns of neuronal firing (Extended Data Fig.11), similar to the reactivation of spatial trajectories in the firing of CA1 place cells during SWRs. Thus, SWRs likely serve as a common mechanism to bind together a diverse array of spatial, sensory and temporal information to consolidate spatial and non-spatial declarative memories, both in rodents and humans^15,16,17^.

## METHODS

### Surgical Procedures

Adult male mice (C57Bl6/J background, ~3-months old) were kept in the vivarium on a 12-hour light/dark cycle and were house 3-4 per cage before surgery and individually after it. All experiments were approved by the Institutional Animal Care and Use Committee at Columbia University. We used transgenic mice to obtain selective expression of cre-dependent virally expressed opsines in the hippocampal subregions CA2 (Amigo2-Cre animals), CA3 (Grik4 animals) or CA1 (Lypd1 animals) for optogenetic manipulations.

In a first surgical procedure, animals were injected with DIO-ChR2 or or DIO-eArch3.0. Mice were anesthetized with isoflurane and analgesic drugs were provided. A craniotomy was performed under stereotaxic guidance in the corresponding coordinates over the target region. Using a Nanoject II (Drummond Scientific), a glass pipette previously loaded with the corresponding virus was then inserted until final depth. Two injections (one per hemisphere) of 200 nL each were performed. After injections, the skin was sutured and treated with broad spectrum antibiotic. Animals were left in the vivarium for recovery for ~3 weeks.

In a second surgical procedure, animals were implanted with silicon probes (−1.94 antero-posterior, −2.2 medio-lateral from Bregma) and optic fibers (AP −1.94, ML +−2.2) to record local field potentials (LFP) and spikes from the dorsal hippocampus and stimulate or just with optic fibers, bilaterally, to silence a specific subset of cells during behavior, similar to previously described procedures^9,10,36^. The probes (5 shanks × 12 electrode sites -A5×12-Poly2-5mm-20s-160- or 4 shanks × 16 electrode sites-A4×16-Poly2-5mm-20s-160-) were mounted on custom-made micro-drives to allow their precise vertical movement after implants. A fiber optic was previously attached under microscopic guidance in one of the shanks of the silicon probe. Probes were inserted above the target region and the micro-drives were attached to the skull with dental cement. In the contralateral hemisphere, a fiber optic (200 um) was implanted over the final region and cemented to the skull. For the animals implanted just with fiber optics, both fibers (one per hemisphere) were implanted in the final position and attached to the skull with dental cement. Craniotomies were sealed with sterile wax. A stainless-steel wire was inserted over the cerebellum to serve as ground and reference for the recordings. Finally, a copper mesh was attached to the skull and secured with metal bars to strengthen the implant. The copper mesh was attached to the skull with dental cement and connected to the ground wire and electrodes’ ground and reference wires to attenuate the contamination of the recordings by environmental electrical noise. After post-surgery recovery (~1week), probes were lowered gradually in 75 to 150 um steps per day until the desired position was reached. We used physiological landmarks and characteristic LFP patterns^36,37^ to identify the pyramidal layers of the different subregions of the hippocampus (based in current source density analysis or independent component analysis)^38,39^ and optogenetic responses.

### Optogenetic experiments

For real-time SWRs manipulations, 100 or 200 um core multi-mode optic fibers (Thor Labs) were collimated to a blue light (475 nm) emitting laser diode (PL-450, Osram). The laser diodes were later connected to current source stimulators (Thor Labs) using a light cable to facilitate the free movement of the animal^9,24,25,40^. For the stimulation session, each of these two diodes were connected to one of the bilaterally implanted fibers.

#### 1. SWRs disruption

A closed-loop system was used in order to detect SWRs online and trigger light stimulation. For SWRs disruption, once a ripple was detected, a short duration (~10 ms) pulse of large amplitude (5-10 mW) was delivered through both fibers to activate ChR2^9^. The intensity was manually adjusted in each animal during test session in the home cage by gradually increasing light power until a population spike was evoked, truncating the ongoing ripple and producing a strong inhibition of pyramidal cell firing, similar to the effect of electrical stimulation. Pulses were delivered every time a ripple was detected while the animal was in the home-cage, between the learning and the recall trial. Two control stimulation experiments were performed: in the first group, the same cre+ animals were used and, in separate sessions, the same stimulation was delivered upon SWRs detection plus a random delay (500-100ms); in a separate cohort of cre-animals (injected previously with the same virus as cre+ animals), the same stimulation was delivered upon SWRs detection.

#### 2. SWR generation

For SWRs generation, 60 ms, small amplitude (1-2mW) light pulses were used to activate ChR2^9,24,25^. The intensity was manually adjusted in each animal during a test session in the home cage by gradually increasing light power until a ripple was produced. To minimize the broad-band artifact at the onset of the stimulation, trapezoid pulses with ~20ms initial ramp were used. Pulses were delivered constantly at a rate that approximate the spontaneous occurrence of SWRs (0.3 Hz).

#### 3. Pyramidal cell silencing during sleep

For the SWRs silencing experiments^40^, a yellow laser (~556 nm, Opto Engine LLC) was used to deliver the light and activate Arch. 3-5mW light pulses were delivered continuously during 30s every 2 minutes.

#### 4. Pyramidal cell silencing during social interaction

For real-time behavioral manipulations, we streamed online (https://github.com/wonkoderverstaendige/Spotter) the position of the animals and yellow light pulses were delivered continuously when the animals crossed the area of interest to activate Arch^40^.

### Behavioral recordings

After surgery, animals where handled daily and accommodated to the experimenter, recording room, cable and recording arena for 1 week before the start of the experiments. One recording session was conducted per day, with minimum 2 days of separation between successive sessions. In a typical session we recorded neural activity during: 1 h in the home cage, 5 min during exposure to the empty arena, 1 h in home cage, 5 min exposure to the cups, 1 h in home cage (pre-sleep), 5 min exposure to two novel stimulus animals (S_1_ and S_2_) in learning trial 1, 5 min exposure to the same stimulus mice in reversed positions in learning trial 2, 1 h in home cage (post-sleep, where SWRs manipulations were performed in some sessions), followed by a 5-min recall trial where one of the two stimulus mice (S_1_ or S_2_, chosen at random) was presented together with a novel stimulus animal (N). For the 24 h recall experiment, the recall test was conducted on the next day. For the stimulus animals, adult male mice (C57Bl6/J background, ~3-months old) were used.

The position of the animal was tracked online using two PCB mounted LED’s (blue, red) and an overhead camera (Basler, 60 Hz). Offline tracking of the animal was validated using pose estimation user-defined features with a deep learning algorithm (https://github.com/AlexEMG/DeepLabCut)^41^. In each trial we measured the total time the subject mouse spent exploring the two stimulus mice (t_S1_, t_S2_) in the two learning trials and the differential time spent in social exploration of the stimulus mouse (e.g. t_S1_) compared to the novel mouse (t_N_) in the recall trial. The differential interaction was quantified by a “Discrimination Index” = (t_N_ – t_S1_)/(t_N_ + t_S1_). Social exploration was defined as time spent inside the interaction zone (considered as 5 cm around the cup location).

Electrophysiological recordings were conducted using Intan RHD2000 interface board (http://intantech.com/RHD2000_evaluation_system.html) at 20KHz.

### Real time SWR detection

Two LFP channels were used for real-time SWRs detection^9,25^. The ripple was detected in one channel of the CA2 region, previously identified by optogenetic tagging of cells response. LFP was filtered between 100-300 Hz. A second channel was used for noise detection, selected from the neocortex where no ripple activity was present and filtered in the same frequency band. The root-mean square (RMS) of the two signals was then computed by a custom-made analog circuit. Signals were fed into a data acquisition interface for real-time processing of LFP channels by a programmable processor (Cambridge Electronic Design – CED) at 20kHz sampling rate. Amplitude thresholds for ripple and noise were manually adjusted for each animal before the start of each recording session based on physiological features. SWRs were defined as events crossing the ripple detection but not crossing the noise signal detection^9,25^.

### Tissue Processing and Immunohistochemistry

Mice were transcardially perfused using saline then 4% PFA in PBS. The brains were quickly extracted and incubated in 4% PFA overnight. After 1 h washing in 0.3% glycine in PBS, 60-μm sections were prepared using a Leica VT1000S vibratome. Sections were permeabilized and blocked for 2 h at room temperature in PBS containing 5% goat-serum and 0.5% Triton-X. Sections were incubated overnight at 4 °C with primary antibodies chicken anti-GFP (1:1,000, AVES Labs, Cat# GFP-1020, RRID: AB_10000240) and rabbit anti-PCP4 (1:200, Sigma-Aldrich, Cat# HPA005792, RRID: AB_1855086), diluted in PBS containing 5% goat-serum and 0.1% Triton-X in PBS. The sections were washed three times for 15 min in PBS at room temperature and secondary antibodies were applied overnight at 4°C in PBS containing 5% goat-serum and 0.1% Triton-X. All secondary antibodies were raised in goat and purchased from ThermoFisher Scientific. The secondary incubation was performed with anti-chicken conjugated to Alexa 488 (Cat# A11039, RRID: AB_142924) and anti-rabbit conjugated to Alexa 647 (Cat# A21244, RRID: AB_2535812). Slices Hoechst 33258 (ThermoFisher Scientific, Cat# H21491) staining was applied at 1:1,000 for 10 min in PBS at room temperature before mounting the section using fluoromount (Sigma-Aldricht). An inverted confocal microscope (Leica, LSM 700) was used for fluorescent imaging while an epifluorescence microscope (Olympus Bx61VS) was used to verify the tracks of electrodes and optical fibers.

### Quantification and statistical analysis

#### Spike Sorting and Unit Classification

Neuronal spikes were detected from the digitally high-pass filtered LFP (0.5-5kHz) by a threshold crossing-based algorithm (Spikedetekt2). Detected spikes were semi-automatically sorted using ‘Kilosort’ (https://github.com/cortex-lab/KiloSort), followed by manual curation of the clusters using ‘Phy’ (https://github.com/kwikteam/phy)^42^. Autocorrelograms and waveform characteristics assisted by monosynaptic excitatory and inhibitory interactions were used to select and characterized well-isolated units and separate them into pyramidal cell and interneuron classes^43^. Only well isolated single units were included in all analysis.

#### Offline Detection of SWRs

To detect ripples, the wide-band signal was band-pass filtered (difference-of-Gaussians; zero-lag linear phase FIR), and instantaneous power was computed by clipping at 4 SDs, rectified and low-pass filtered. The low-pass filter cut-off was at a frequency corresponding to p cycles of the mean band-pass (for 80-250 Hz band-pass, the low-pass was 55 Hz). The mean and SD of baseline of LFP were computed based on the power during non-REM sleep. Subsequently, the power of the non-clipped signal was computed, and all events exceeding 4 SDs from the mean were detected. Events were then expanded until the (non-clipped) power fell below 1 SD; short events (<15ms were discarded)^9,10,24,25^.

#### Ripple spike content analysis

Well-isolated putative units with at least 100 spikes in a given session were included in the analysis. For all individual units, spikes in a [−300, +300] ms peri-ripple or stimulus onset window were collected and firing rate histograms (1ms time bin) were constructed. All histograms for the same subregions and same type of units (pyramidal cells) were pooled. The firing rate histograms were z–scored and smoothed using a Gaussian kernel (SD = 5ms). Only spikes within the interval of the detected ripples were considered for the ripple content analysis. In-ripple firing rate was calculated as number of spikes divided by ripple duration and averaged for all events. The probability of participation of individual units in ripples was defined as the number of events in which a neuron fired at least one spike during the ripple divided by the total number of ripples detected. Within-ripple gain was calculated as in-ripple firing rate divided by the baseline firing rate of the cell^9,24,25^.

#### Reactivation Analysis

Explained variance (EV) and reverse explained variance (REV) were calculated in each session using the cell pairs of interest. EV and REV were calculated for pyramidal cells only using a previously described approach^21,22^. EV is defined as:

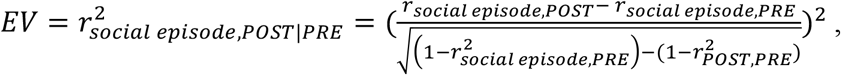

where *r_social episode,POST_* and *r*_social episode,PRE_ are the Pearson’s correlation coefficients of pairwise correlation matrices between the social learning episodes and SWRs times of the post-sleep and pre-sleep periods, respectively, and r_*POST,PRE*_ is the Pearson’s correlation coefficient of the pairwise correlation matrices between the post-sleep and pre-sleep periods. For REM sleep, non-REM sleep or SWR times EV and REV measurements, all epochs/events were detected individually and pooled together in both pre- and post-sleep periods. Pairwise neuronal correlations for EV and REV were calculated using the Pearson correlation coefficient on 100 ms binned spike trains. The coefficients were calculated for pre-sleep, task, and post-sleep periods separately and correlation matrices were then computed. The correlations between all pairwise combinations were calculated and used to assess the percentage of variance in the post sleep period that could be explained by correlation values established during the task trials, while controlling for pre-existing correlations in the pre-sleep session. The control value (REV) was obtained by switching the temporal order of the pre- and post-sleep sessions. For assessing the relative contribution to the explained variance of a single pair of neurons, the EV was calculated for all the pairs without including the pair of interest. Subtracting this value from the total EV value, a relative contribution for the corresponding pair of neurons was calculated. To calculate the average contribution of each neuron to the EV, the relative contribution for all pairs in which a particular neuron participated was averaged^22^.

For detecting cell assembly patterns, we used an unsupervised statistical framework based on a hybrid principal component analysis (PCA) followed by independent component analysis (ICA) as previously reported^20,23^. In brief, spike trains of each neuron were binned in 20-ms intervals for the whole session and z-scored firing rates were calculated for each bin. Spike trains were convolved with a Gaussian kernel (SD = 10 ms) and the matrix of firing correlation coefficients for all pairs of neurons constructed. Next, we calculated the number of assemblies based on those principal components whose eigenvalues exceeded the threshold for random firing correlations (Marčenko-Pastur distribution). Independent component analysis was then used to determine for each assembly (component) the vector of weights with which each neuron’s firing contributes to that assembly. To measure the strength of each assembly’s activation as a function of time in a given episode, we multiplied the convolved z-scored firing rate of a given neuron at a given time by the weight of that neuron’s contribution to a given assembly. We then summed the product of these weighted spike counts for all non-identical pairs of neurons to provide the assembly activation strength at the given time point. Assembly activity was only considered when its expression strength exceeded a threshold of 5^20^. Reactivation strength of an assembly was defined as the difference in its average expression strength during post-sleep minus that during pre-sleep. Larger reactivation numbers indicate a greater social-learning-dependent increase in assembly activation. Assembly activation was assessed inside and outside the two social interaction areas surrounding the stimulus mice. Discriminant assemblies were defined as assemblies that were activated significantly more strongly during the interaction with one of the two stimulus mice compared to the other in the two social learning trials (p < 0.05, Student’s t-test).

#### Place cells and cell analysis

Spiking data was binned into 1-cm wide segments of the camera field projected back onto the maze floor, generating raw maps of spike counts and occupancy probability. A Gaussian kernel (SD = 5 cm) was applied to both raw maps of spike and occupancy, and a smoothed rate map was constructed by dividing the smoothed spike map by the smoothed occupancy map. A place field was defined as a continuous region, of at least 15 cm^2^, where the firing rate was above 10% of the peak rate in the maze, the peak firing rate was > 2 Hz and the spatial coherence was > 0.7. Spatial correlation between trials was calculated as the average pixel-by-pixel correlation of the smoothed firing rate maps of each cell. The rotated spatial correlation was calculated in the same way but rotating 180^0^ the rate map in trial 2, so the position of mice S1 and S2 matched that of trial 1. The distance of each place field to the cups was calculated from the center of mass of the field to the center of the closest cup. The spatial information content and selectivity of rate maps were calculated following previously described methods^44,45^. Information index per spike (bits/spike) represents the mean firing rate of the cell at location X, multiplied by the probability of the animal of being at location X and the binary logarithm of the mean firing rate of the cell at location X divided by the overall firing rate of the cell. The selectivity index represents the maximum firing rate of the rate map (spikes divided by time spent in each bin of 5cm) divided by the mean firing rate.

For the classification of spatial and social cells, k-means clustering was performed with all the normal and rotated correlation values for all pyramidal cells active during the trials of the task. The ‘cityblock’ metric was used as distance measure. We found the optimal number of clusters in the data to be five by employing the silhouette criterion.

#### Statistical Analysis

All statistical analyses were performed with MATLAB functions or custom-made scripts following previously described methods^1–10,20–26^. The unit of analysis was typically identified single cells, assemblies of cells or SWRs events. In some cases, the unit of analysis was sessions or animals, and this is stated in the text. Unless otherwise noted, for all tests, non-parametric two-tailed Wilcoxon rank-sum (equivalent to Mann-Whitney U-test) or Wilcoxson signed-rank were used. For multiple comparisons Tukey’s honesty post hoc test was employed. On box plots, the central mark indicates the median, bottom and top edges of the box indicate 25th and 75th percentiles respectively, and whiskers extend to the most extreme data points without considering outliers, which were also included in statistical analyses. Due to experimental design constraints, the experimenter was not blind to the manipulation performed during the experiments (i.e., optogenetic manipulations).

## DATA AND SOFTWARE AVAILABILITY

Datasets and analytical tools included in this study will be made available upon request.

## ACKNOWLEDGMENTS

We thank Gyorgy Buzsaki for insightful comments and resource sharing throughout the course of the project, members of the Siegelbaum lab for insightful comments and discussions of the manuscript and Torcato Meira for help with designing the behavioral paradigm.

## AUTHOR CONTRIBUTIONS

Conceptualization: A.O. and S.A.S. Experiments and data collection: A.O. Data analysis: A.O and A.F.R. Immunohistochemistry: F.L. Writing – original draft: A.O. Writing – review and editing: A.O. and S.A.S. Figures creation: A.O. Supervision: S.A.S. Funding acquisition: S.A.S.

## CORRESPONDING AUTHORS

Steven A. Siegelbaum; Azahara Oliva

## EXTENDED DATA FIGURES AND LEGENDS

**Supplemental Figure 1:**
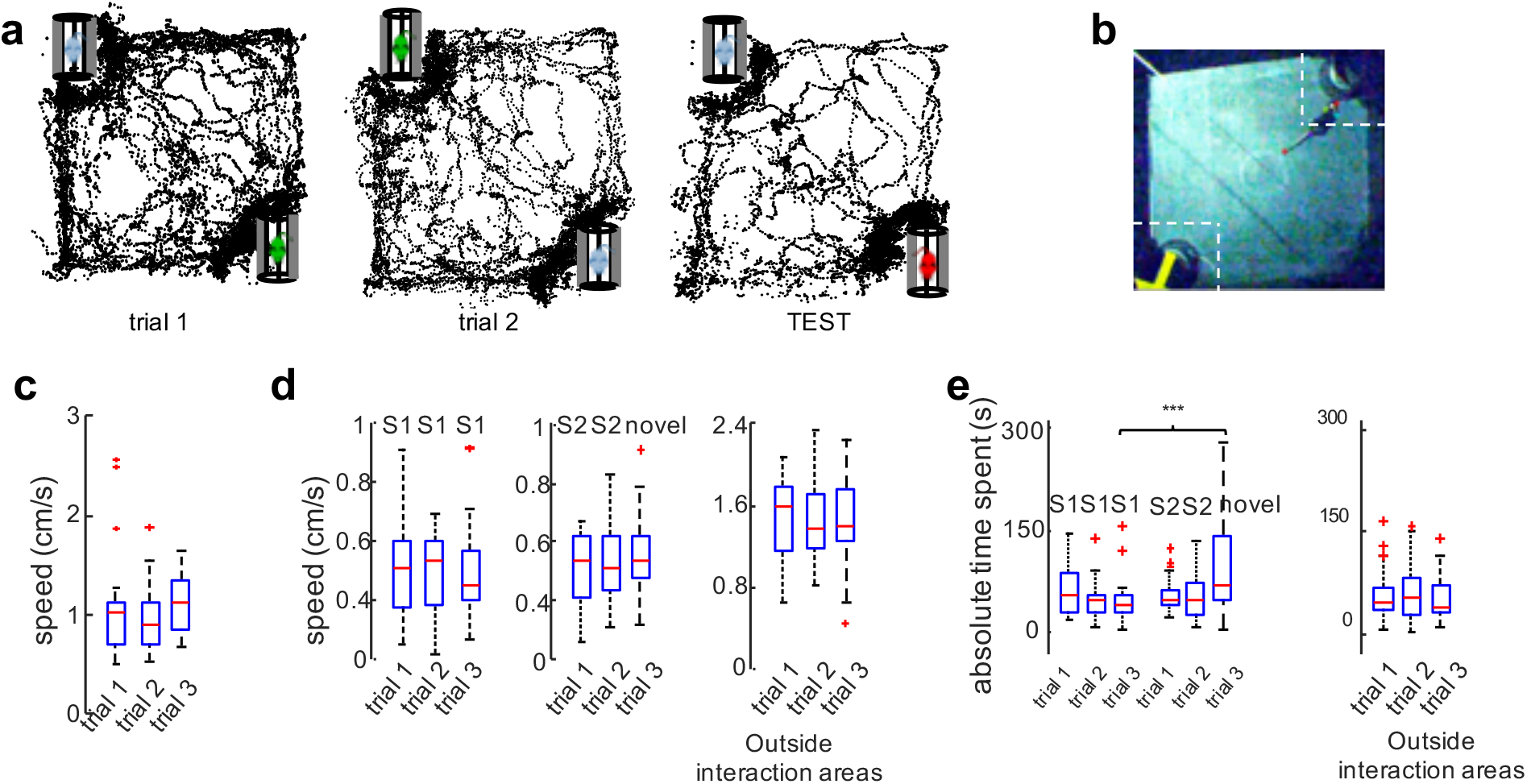
Behavioral Features during social memory task. **a**) Representative animal trajectories during the task show higher time spent around the novel animal (red) in the test trial. **b**) Example video frame showing pose estimation calculated with DeepLabCut (color markers). Interaction zones were defined as 10 cm by 10 cm squares in the two corners where the cups were located. **c**) Average speed of the animals was not different among trials (one-way ANOVA: F(2) = 0.92, P > 0.05). **d**) Average speed inside interaction zones around S1 (left plot), S2 or novel mouse (middle plot) did not differ among trials (one-way ANOVA: F(2) = 3.14, P > 0.05). Average speed did not differ outside interaction zones (one-way ANOVA: F(2) = 1.58, P > 0.05). **e**) Total time spent inside the interaction zone around S1, S2 or novel mouse (left plot). Total time interacting with novel mouse during recall trial was greater than with either familiar mouse in any other trial (p < 0.01 for novel versus S1 interaction during third trial and p < 0.05 for novel versus S1 interaction during all of the other trials, two-way ANOVA mouse × trial followed by Tukey post hoc test for multiple comparisons). Total time spent outside interaction zones (right plot) was not different between trials (one-way ANOVA: F(2) = 0.58, P>0.05).

**Supplemental Figure 2:**
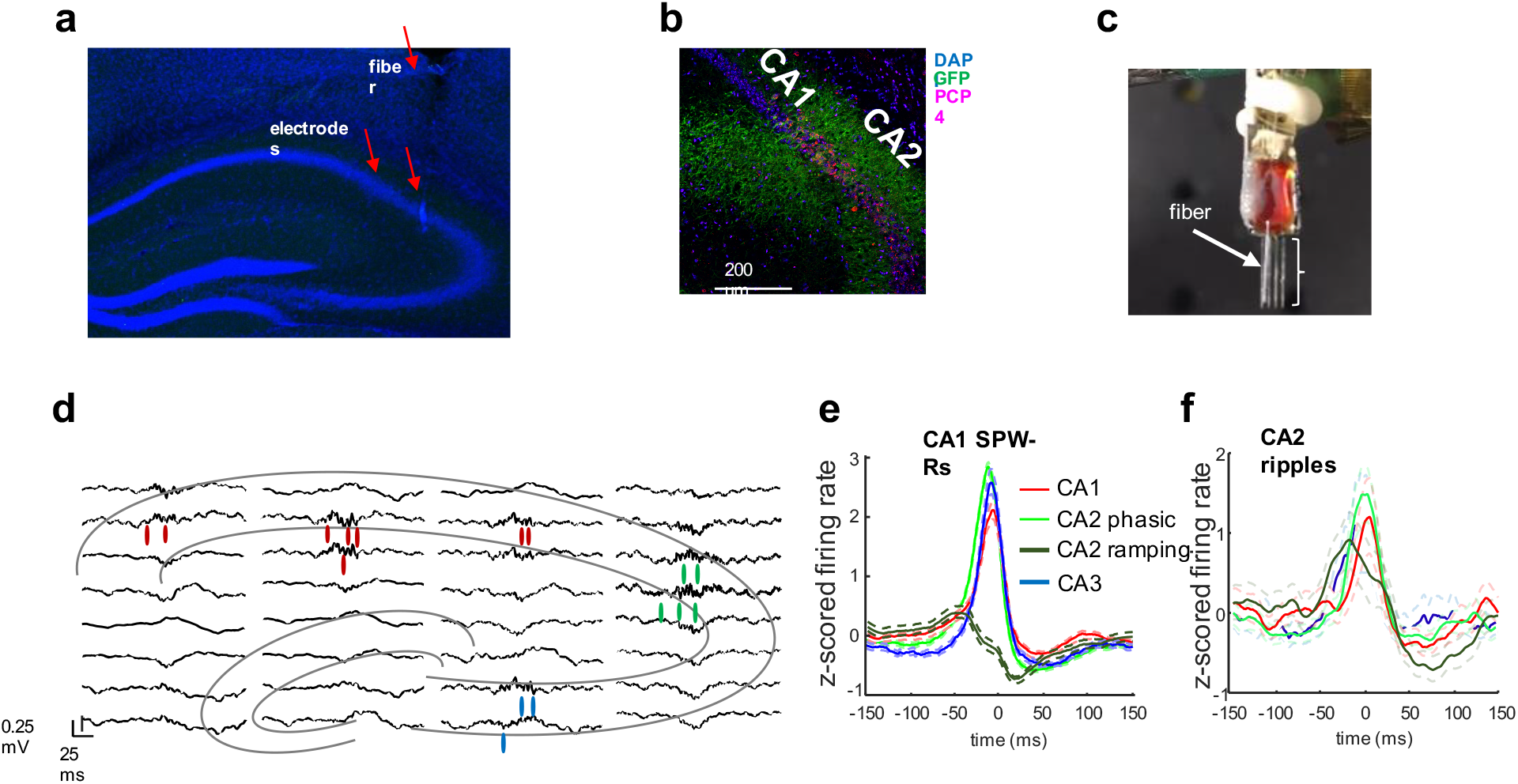
Multi-region electrophysiological recordings. **a**) Representative histology showing electrode tracks spanning CA1, CA2 and CA3 areas, with optic fiber over CA2 pyramidal layer. Blue: DAPI staining. **b**) ChR2-expression in CA2. Blue: DAPI; green: GFP; red: PCP4. **c**) Silicon probe with 100 μm optic fiber glued to one electrode shanks mounted in a movable microdrive to allow for precise localization of the target area. **d**) Representative sample recordings of local field potentials (one trace per electrode) and single units (colored lines show spikes) in several regions of the hippocampus. Each column represents one electrode shank. Approximate location of pyramidal and granular layers is depicted in superimposed outline of hippocampus. **e**) Average firing response curves of single cells from different regions aligned to SWRs detected in CA1. Note that CA2 cells fired before CA1 and CA3 and that a subpopulation of CA2 units (‘CA2 ramping’) become silent upon SWR onset. **f**) Same as in E but firing responses were aligned to ripples detected in CA2. Note that CA2 cells are strongly active during CA2 ripples. These results replicate a previous report in rats^21^.

**Supplemental Figure 3:**
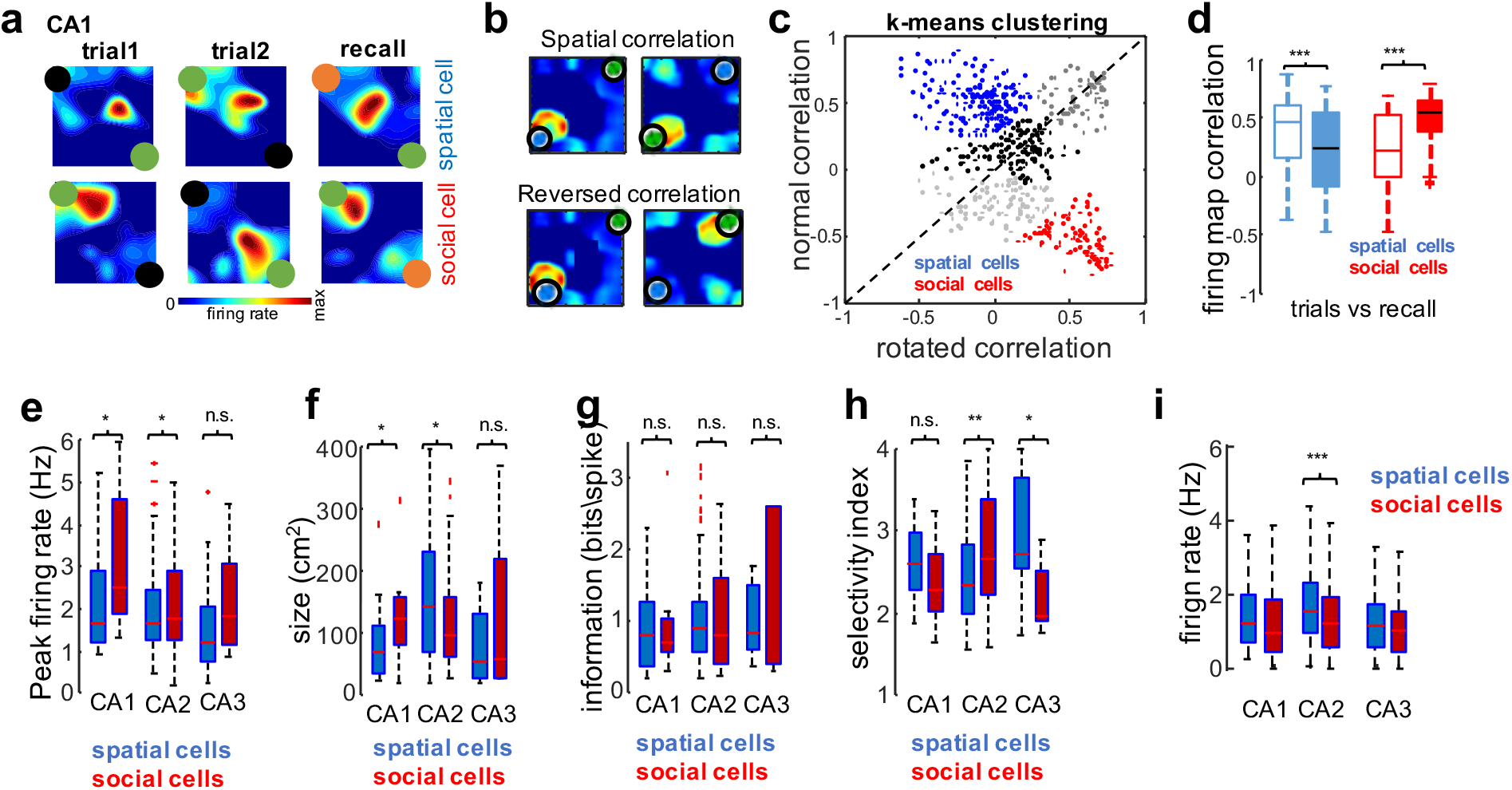
Single cell firing properties during social memory task. **a**) Example CA1 firing maps for a spatial cell (top; standard correlation=0.78, rotated correlation=0.12 for trial 1 compared to trial 2) and social cell (bottom; standard correlation = −0.64, rotated correlation = 0.67). **b**) Standard spatial correlation was computed as averaged pixel-wise correlation of the firing maps from trial 1 and 2 (top). Rotated correlation was calculated after rotating the map for trial 2 180 degrees (bottom). **c**) K-means clustering of normal and rotated correlation values for all cells resulted in 5 clusters. One corresponded to spatial cells (blue), with high normal and negative rotated correlations. Another corresponded to social cells (red), with high rotated and negative normal correlations. **d**) We checked that the spatial correlation of firing maps for social and spatial cells demonstrated between learning trials was also preserved in subsequent recall. For that, we compared the recall trial with the learning trial in which the same familiar mouse was in the opposite corner. Spatial cells had higher standard than rotated correlations in learning compared to recall trials (p < 10^−4^, sign-rank test), evidencing the stability of the allocentric spatial representation. Social cells, on the other hand, had higher rotated than standard correlations (p < 10^−7^, sign-rank test), indicating that firing maps remapped with the position of the familiar conspecific. **e-h**) Properties of spatial and social cells in the three regions: (**e**) mean in-field firing rate; (**f**) field size; (**g**) spatial information in bits per spike (see methods); (**h**) selectivity index (see methods). **i**) Session-wide average firing rate for social and spatial cells from the different subregions. */**/*** p < 0.05/0.01/0.001, rank-sum test. **j**) Firing rate of pyramidal cells during running and immobility. */** p < 0.05/0.01, rank-sum test.

**Supplemental Figure 4:**
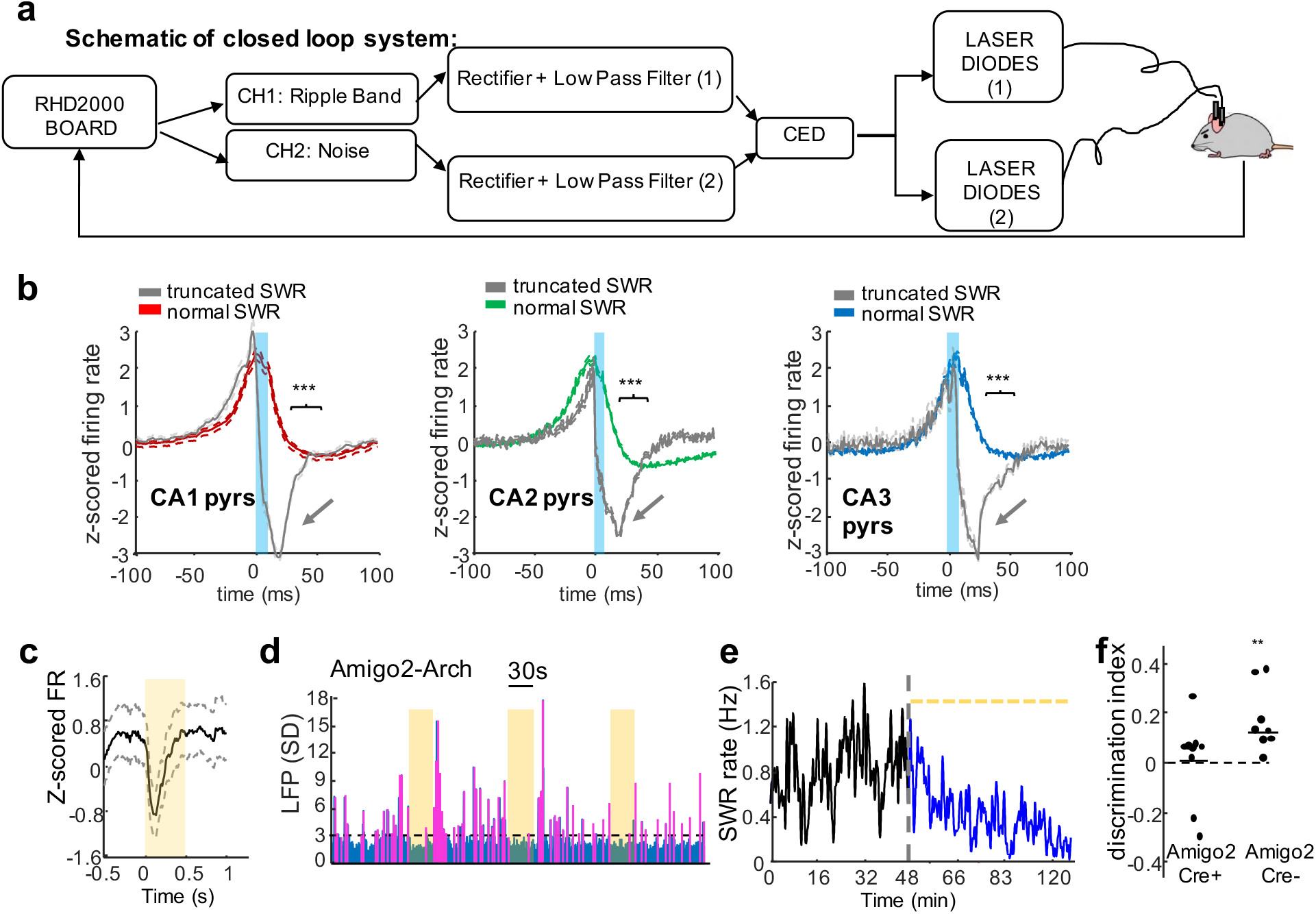
Reactivation of hippocampal cells during SWRs. **a**) Schematic of closed-loop SWR truncation system. **b**) CA1, CA2 and CA3 average firing responses to normal and truncated SWRs show strong suppression of firing after light stimulation (blue rectangle) (n = 53, 148, 87 CA1, CA2, CA3 cells; p < 10^−6^, = 9.7×10^−7^, = 1.14×10^−8^, respectively sign-rank test). Firing of CA2 pyramidal cells was suppressed by brief yellow-light pulses (yellow rectangle) in Amigo2-Cre animals expressing eArch3.0-DIO-eYFP in CA2. Curves show mean and SEM (n = 58, p < 0.03, sign-rank test). **d**) Example session in which 30-s pulses of yellow light (yellow rectangles) were delivered once every two minutes to CA2 of Amigo2-Cre mice expressing Arch3.0. Blue trace is ripple-band (100-200 Hz) power in the CA1 pyramidal layer and magenta detected SWRs. Note suppression of SWRs during illumination. **e**) Example session showing SWR rate decrease due to the stimulation (p < 0.0246, Wilcoxon rank sum test). Dashed line shows time of light stimulation sand black and blue traces show SWR rate before and during period of photostimulation, respectively. **f**) Social memory recall was impaired by light pulses during post-sleep period in Amigo2-Cre mice injected with AAV-DIO-Arch (n=9, p>0.05, t-test whether discrimination index differed from 0) but not in Cre-littermates controls that underwent the same protocol (n=8, p<0.01).

**Supplemental Figure 5:**
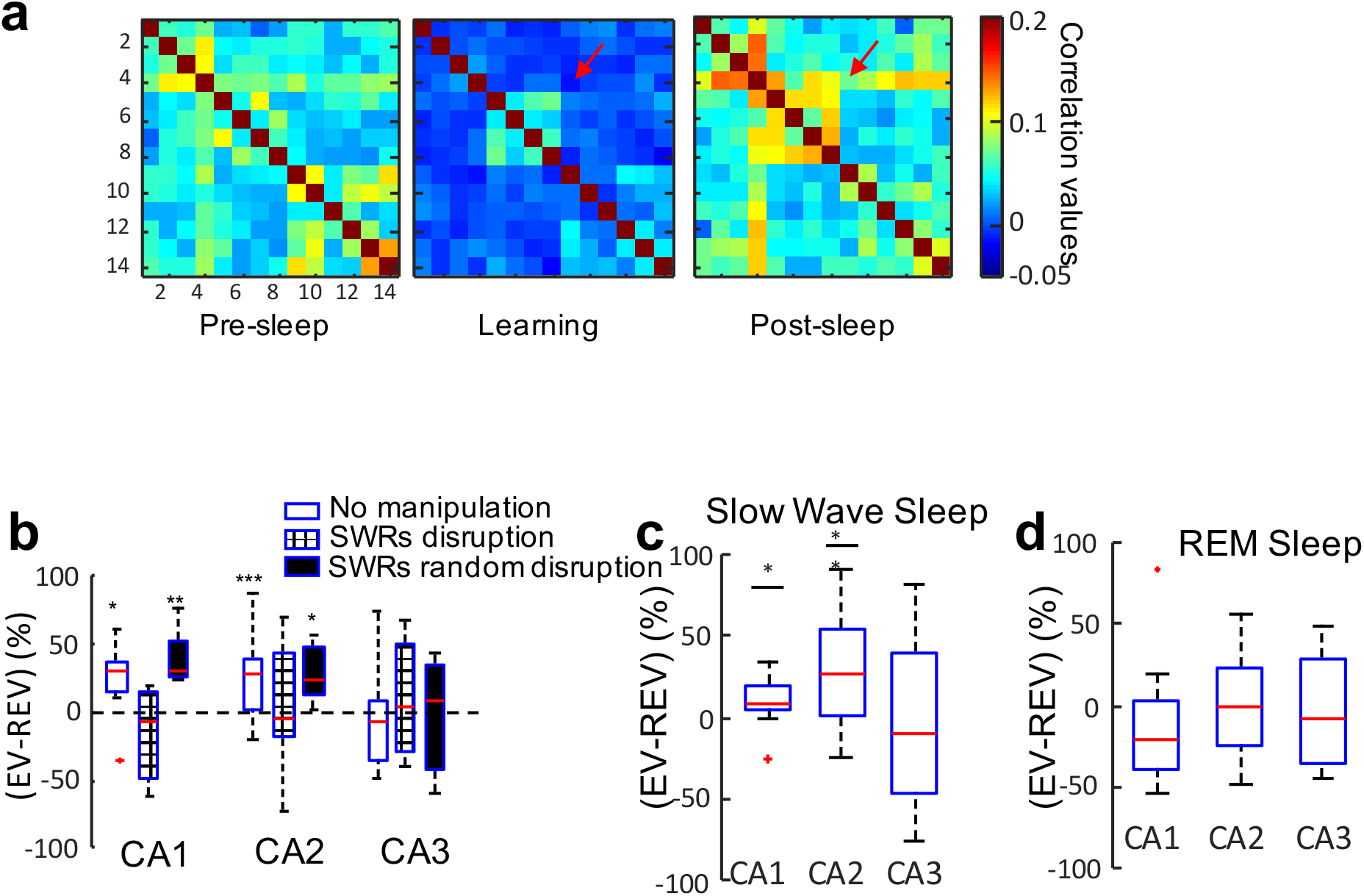
Reactivation of hippocampal cells during SWRs following social learning. **a**) Example of cell pair firing rate correlation matrices for pre-sleep, learning trials and post-sleep. Bar, Color-coded r values. Note increase in post-sleep coactivation in some cell pairs that were coactive during the task (red arrow). **b**)(EV – REV) difference was used to quantify post-sleep reactivation in control sessions (no manipulation), sessions with optogenetic SWR disruption, and sessions with random optogenetic stimulation. Significant reactivation was observed in CA1 and CA2 control sessions (p = 0.03 and 0.0028, respectively, Wilcoxon rank sum test) or following random stimulation (p=0.008 and 0.04 for CA1 and CA2, respectively, Wilcoxon rank sum test). There was no significant reactivation in CA1 or CA2 with SWR disruption. CA3 failed to show significant reactivation in any session (p>0.05). **c**) Significant reactivation (EV-REV difference) was observed in CA1 and CA2 during entire slow-wave sleep period but not during REM sleep period (**d**).

**Supplemental Figure 6:**
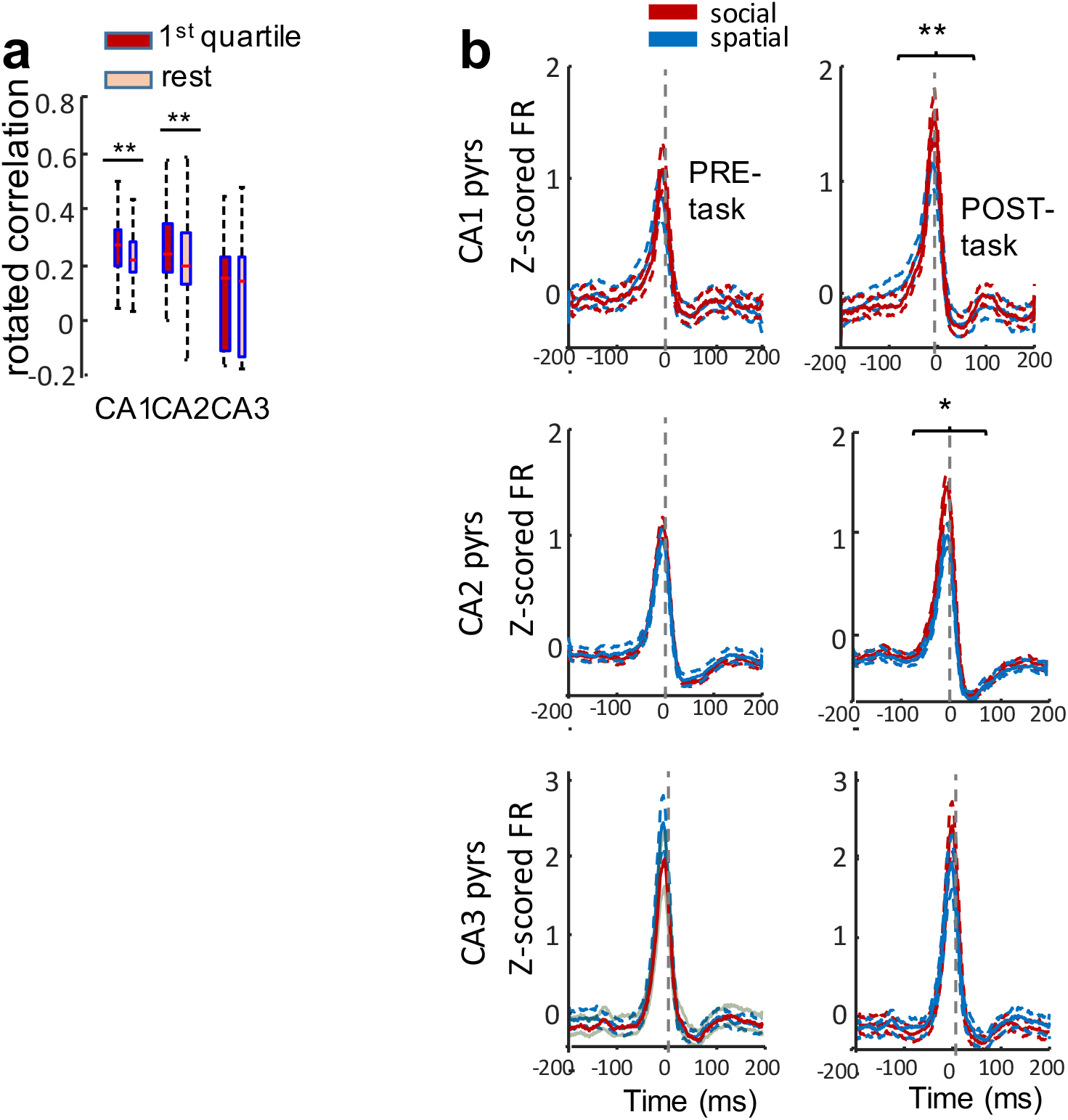
Reactivation of single cells during SWRs in the social memory task. **a)** Cells from CA1 and CA2 that contributed the most to the total explained variance (1^st^ quartile) had a significantly higher rotated spatial correlation than the rest of the cells (p = 0.0354 and p = 0.0223 for CA1 and CA2 respectively; p>0.05 for CA3, Wilcoxon signed rank test). **b)** Average peri-SWR firing rate responses for social and spatial cells from each region in pre- and post-sleep. Note that social cells show higher SWR firing rates in post-sleep but not pre-sleep sessions (CA1: n = 151, p = 0.0055; CA2: n = 306, p = 0.016; CA3: n = 79, p > 0.05)

**Supplemental Figure 7:**
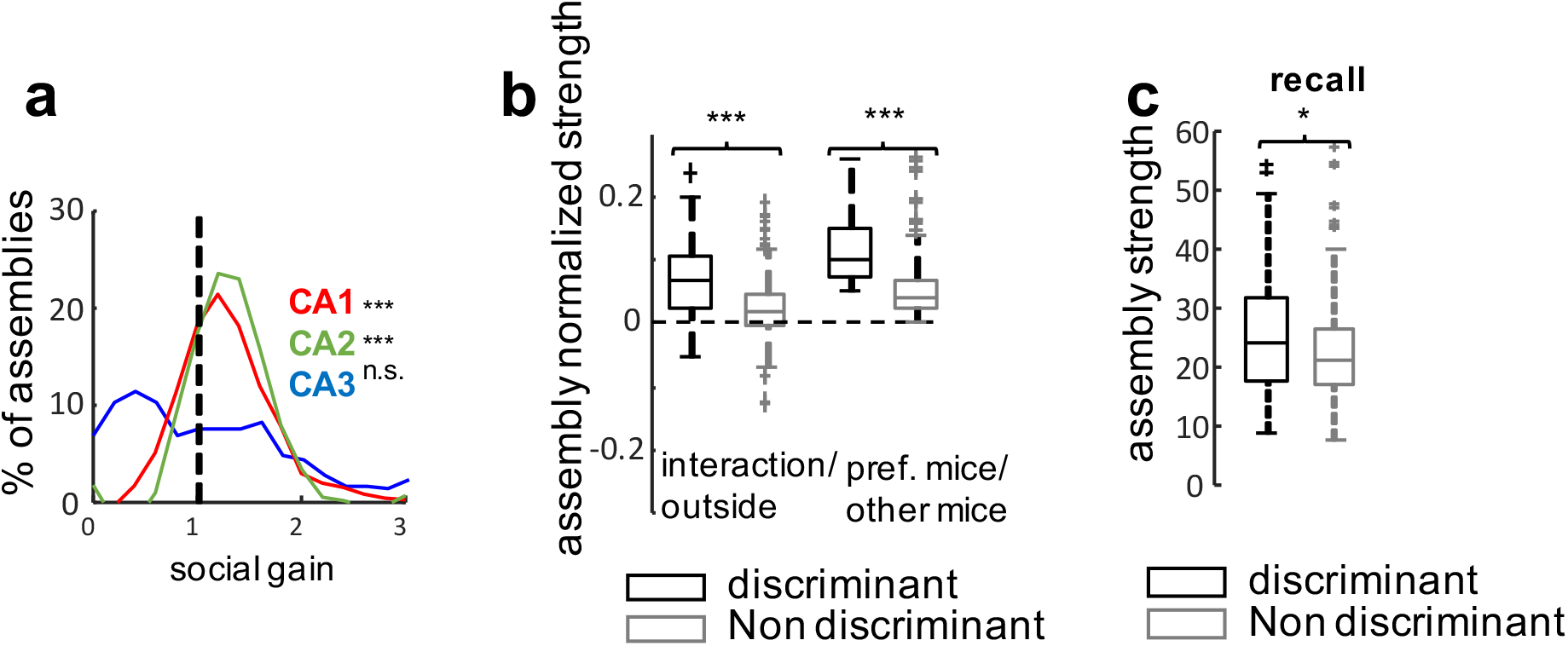
Assembly activity strength during the social memory discrimination task. **a**) Distribution of assembly social gain values from different regions. Assembly social gain defined as mean assembly strength during exploration within interaction zone divided by mean assembly strength during exploration outside interaction zone. Social gain was significantly greater than 1 for CA1 (p < 10^−6^, sign-rank test)) and CA2 (p < 10^−19^) but not CA3 (p > 0.05). **b**) Left pair of bars: normalized social assembly strength (difference between assembly strength inside social interaction zone minus strength outside social interaction zone divided by the sum of the two strengths) was higher for discriminant than non-discriminant assemblies (p < 10^−3^, rank-sum test). Right pair of bars: normalized social discrimination assembly (difference between assembly strength during interaction with preferred mouse minus the strength during the interaction with the other mouse divided by the sum of these two strengths) was greater for discriminant compared to non-discriminant assemblies (p < 10^−8^). **c**) Discriminant assemblies were reactivated during recall trial significantly more strongly than non-discriminant ones (p = 0.0421).

**Supplemental Figure 8:**
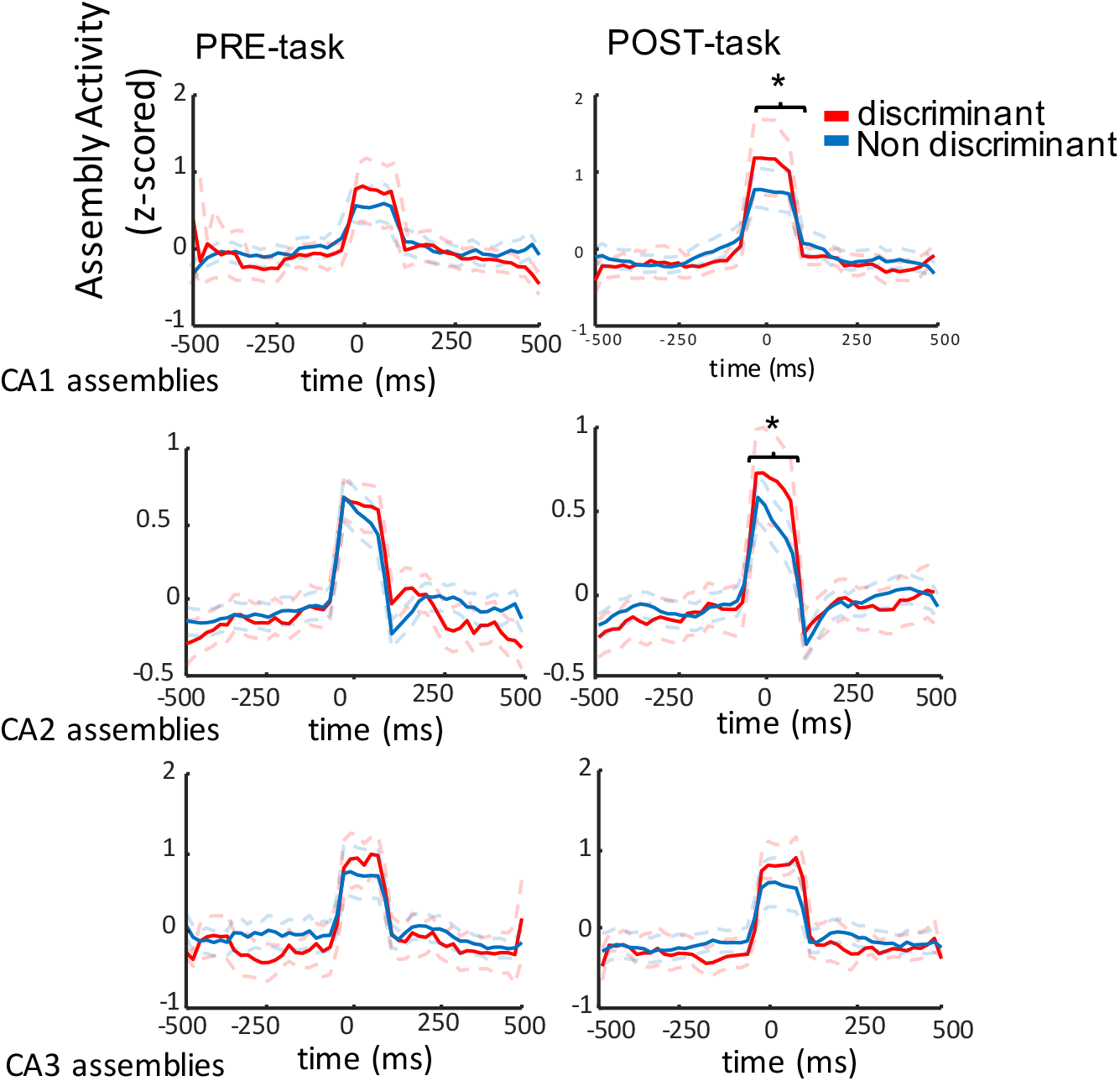
Activation of cell assemblies during SWRs in pre-sleep and post-sleep sessions. Average peri-SWR activation of discriminant and non-discriminant assemblies in different hippocampal regions. CA1: n=116, p=0.0252; CA2: n=213, p =0.0144; CA3: n=59, p>0.05.

**Supplemental Figure 9:**
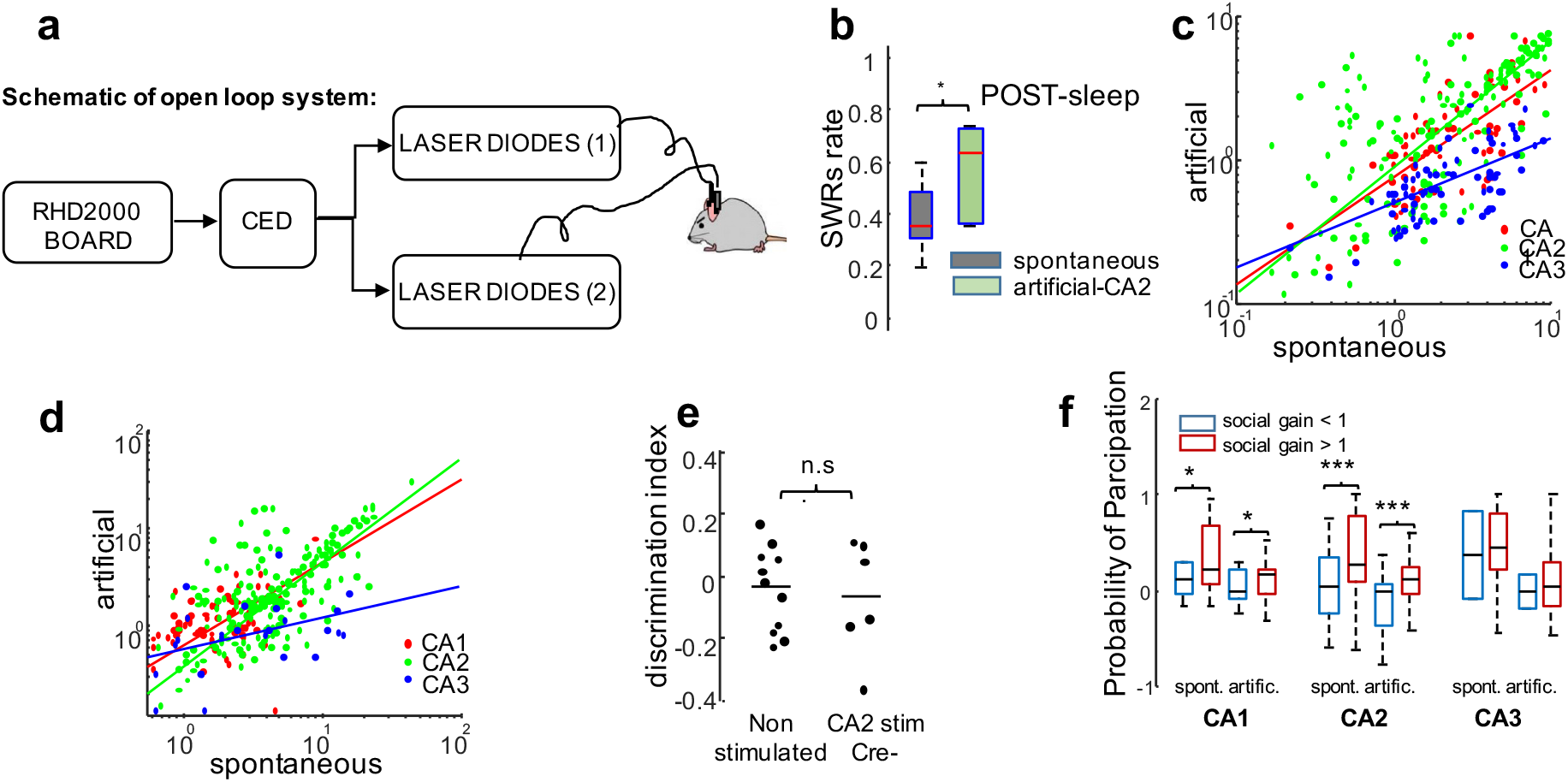
Generation of CA2 ripple oscillations enhances social memory recall. **a**) Schematic of open-loop ripple generation system. **b**) Rate of ripples in stimulated sessions was significantly higher than without artificial ripple generation (p < 0.05, rank-sum test). **c**) Firing rates of all pyramidal cells during spontaneous versus artificially generated CA2 ripples were highly correlated: CA1 (n = 67; r= 0.63, p < 10^−13^, Pearson’s correlation), CA2 (n= 147; r= 0.75, p < 3×10^−22^) and CA3 (n = 40; r = 0.48, p = 0.01). **d**) Firing rate gain (increase firing rate during ripples divided by average firing rate) of pyramidal cells during spontaneous versus artificial ripples for CA1 (n = 67; r= 0.57, p < 10^−6^), CA2 (n= 147; r= 0.74, p < 3×10^−35^) and CA3 (n = 40; r= 0.25, p > 0.05) **e**) Social discrimination index for Amigo2-Cre-littermate controls injected with Cre-dependent ChR2 AAV with and without light stimulation did not differ (p>0.5, n=10). **f**) Effect of social gain on a neuron’s ripple participation gain (post-sleep participation minus pre-sleep participation divided by their sum). CA1 and CA2 cells showed greater ripple participation gain for cells with positive versus negative social gain for both spontaneous SWRs (CA1 and CA2: n = 67 and n = 147; p < 0.05 and p < 3.4×10^−3^, respectively) and artificial SWRs (CA1 and CA2: p < 0.05 and p < 2.8×10^−3^, respectively). CA3 ripple participation gain showed no effect of social gain for either type of SWR (n=40; p>0.05).

**Supplemental Figure 10:**
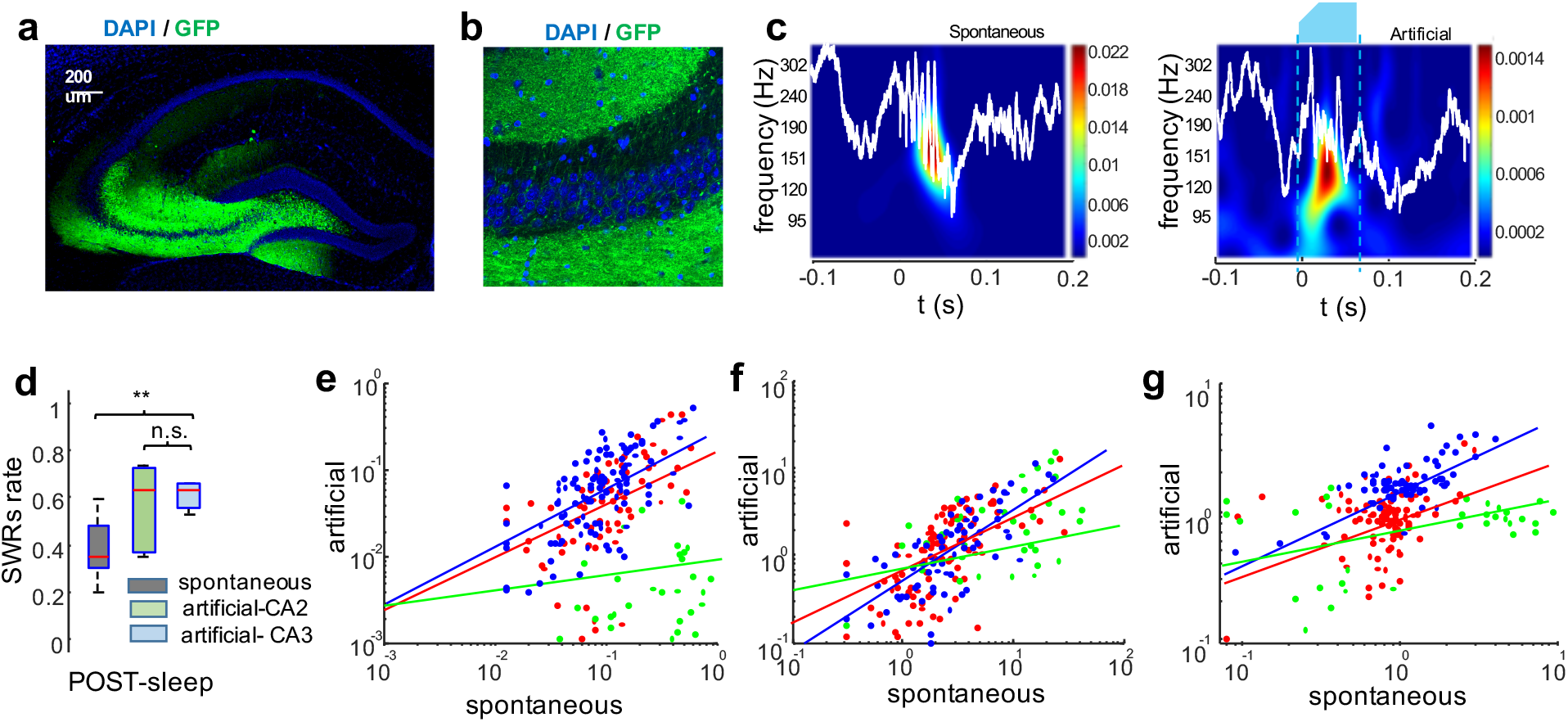
Generation of CA3 ripple oscillations did not extend social memory recall. **a**) Histology of CA3-implanted Grik-4 animals, previously injected with Cre-dependent AAV expressing ChR2-eYFP (green). **b**) Close-up view of the CA3 area. **c**) Example of spontaneous and optogenetically generated ripples in CA3. White lines are LFP from CA2, color maps show wavelet spectrogram, dashed lines indicate period of illumination. **d**) Rate of ripples in sessions with artificial CA3 ripple generation was significantly higher than in non-stimulated sessions (p < 0.003), with no significant difference compared to rate of ripples in response to CA2 stimulation (p > 0.05). **e**) Participation probability (fraction of ripples in which the neuron fire at least one spike) of all pyramidal cells during spontaneous versus artificially generated CA2 ripples were highly correlated: CA1 (n = 96; r= 0.66, p < 7×10^−10^, Pearson’s correlation), CA2 (n= 67; r= 0.34, p < 3×10^−22^) and CA3 (n = 112; r = 0.67, p = 3×10^−15^). **f**) A similar result was obtained comparing firing rates (r= 0.59, 0.49, 0.66; p < 2×10^−7^; 1.7 × 10^−5^; 8×10^−10^; for CA1, CA2 and CA3, respectively) or firing rate gain (**g**) (r= 0.63, 0.47, 0.71; p<1.6×10^−12^, 2.1×10^−3^, 1.7×10^−18^).

**Supplemental Figure 11:**
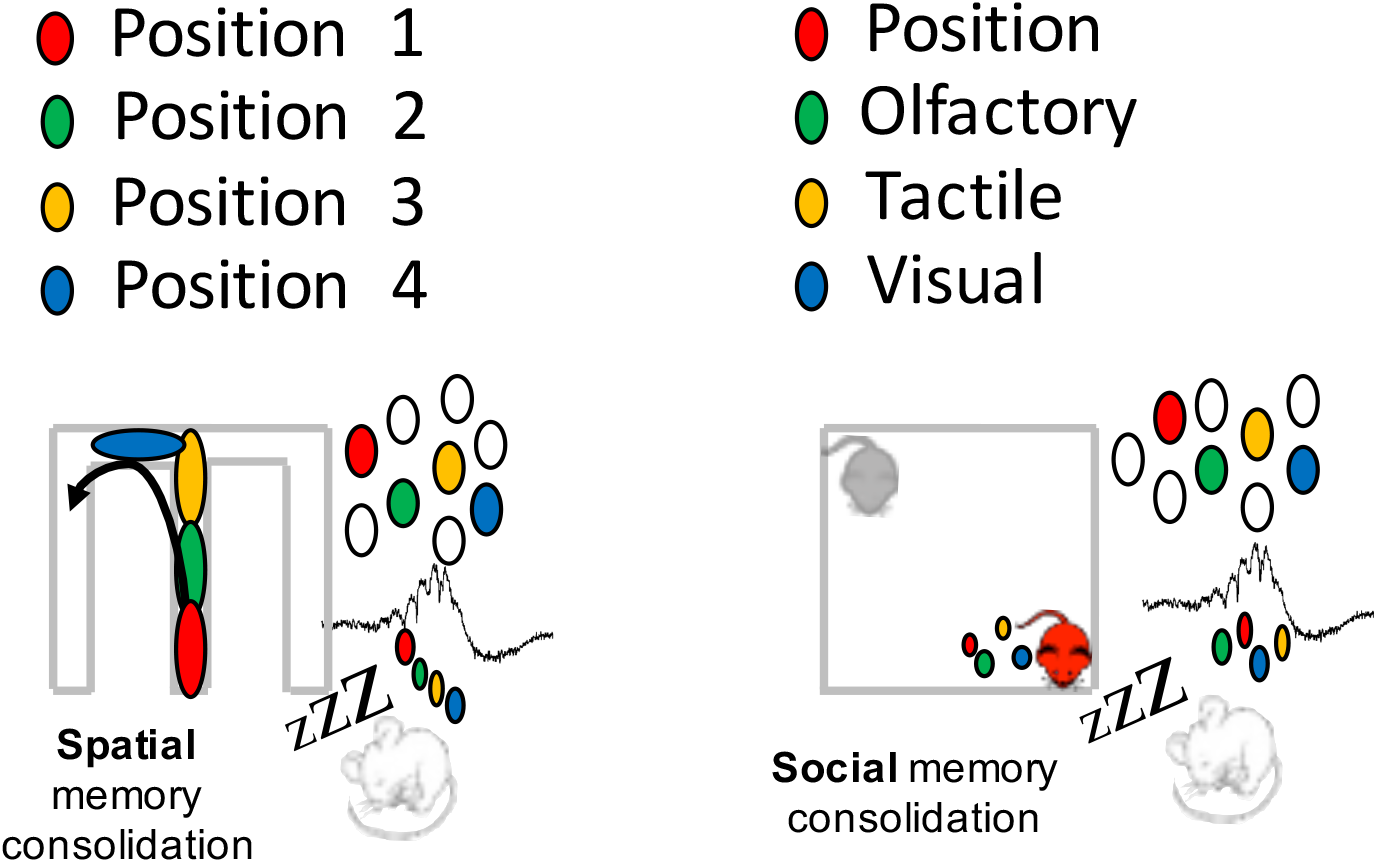
Model of SWRs as integrators of multimodal spatio-temporal-sensory information. **a**) SWRs provide a temporally compressed reactivation of an assembly of place cells that fired during a particular trajectory in a spatial navigation task. b) SWRs reactivate multimodal sensory, spatial and temporal inputs experienced during social exploration to define a compressed “social space”.

## References

1. Wilson M.A. & McNaughton B.L. Reactivation of hippocampal ensemble memories during sleep. Science. 265:676–679 (1994).

2. Diba K. & Buzsáki G. Forward and reverse hippocampal place-cell sequences during ripples. Nat. Neurosci. 10, 1241–1242 (2007).

3. Karlsson M.P. & Frank L.M. Awake replay of remote experience in the hippocampus. Nat. Neurosci. 12, 913–8 (2009).

4. Girardeau G., Benchenane K., Wiener S.I., Buzsaki G. & Zugaro M.B. Selective suppression of hippocampal ripples impairs spatial memory. Nat. Neurosci. 12:1222–1223 (2009).

5. Dupret, D., O’Neill, J., Pleydell-Bouverie, B. & Csicsvari, J. The reorganization and reactivation of hippocampal maps predict spatial memory performance. Nat. Neurosci. 13, 995–1002 (2010).

6. Ego-Stengel V & Wilson MA. Disruption of ripple-associated hippocampal activity during rest impairs spatial learning in the rat. Hippocampus. 20(1),1–10 (2010).

7. Jadhav, S. P., Kemere, C., German, P. W. & Frank, L. M. Awake hippocampal sharp-wave ripples support spatial memory. Science. 336(6087), 1454–1458 (2012).

8. Buzsáki G. Hippocampal sharp wave-ripple: A cognitive biomarker for episodic memory and planning. Hippocampus. 25, 1073–1188 (2015).

9. Fernández-Ruiz A, Oliva A, Fermino de Oliveira E, Rocha-Almeida F, Tingley D & Buzsáki G. Long duration spontaneous and optogenetically prolonged hippocampal sharp-wave ripples improve memory. Science. 364(6445),1082–1086 (2019).

10. Oliva A, Fernández-Ruiz A, Buzsáki G, Berényi A. Role of hippocampal CA2 region in triggering Sharp-Wave Ripples. Neuron. 91(6),1342–1355 (2016).

11. Hitti F.L. & Siegelbaum S.A. The hippocampal CA2 region is essential for social memory. Nature. 508(7494),88–92 (2014).

12. Meira T, Leroy F, Buss EW, Oliva A, Park J & Siegelbaum SA. A hippocampal circuit linking dorsal CA2 to ventral CA1 critical for social memory dynamics. Nat Comm. 9:4163 (2018).

13. Alexander GM, Farris S, Pirone JR, Zheng C, Colgin LL & Dudek SM. Social and novel contexts modify hippocampal CA2 representations of space. Nat Comm. 7:10300 (2016).

14. Eschenko O, Ramadan w, Mölle M, Born J & Sara SJ. Sustained increase in hippocampal sharp-wave ripple activity during slow-wave sleep after learning. Learn Mem. 15(4),222–8 (2008).

15. Norman Y, Yeagle EM, Khuvis S, Harel M, Mehta AD & Malach R. Hippocampal sharp-wave ripples linked to visual episodic recollection in humans. Science. 365(6454), eaax1030. (2019).

16. Vaz AP, Inati SK, Brunel N & Zaghloul KA. Couple ripple oscillations between the medial temporal lobe and neocortex retrieve human memory. Science. 363(6430), 975–978 (2019).

17. Liu Y, Dolan RJ, Kurth-Nelson Z & Behrens T EJ. Human replay spontaneously reorganizes experience. Cell. 178, 640–652 (2019).

18. Alexander GM, Brown LY, Farris S, Lustberg D, Pantazis C, Gloss B, Plummer NW, Jensen P & Dudek SM. CA2 neuronal activity controls hippocampal low gamma and ripple oscillations. eLife. 7:e38052 (2018).

19. Boehringer R, Polygalov D, Huang AJY, Middleton SJ, Robert V, Wintzer ME, Piskorowski RA, Chevaleyre V & McHugh TJ. Chronic loss of CA2 transmission leads to hippocampal hyperexcitability. Neuron. 94, 642–655 (2017).

20. van de Ven GM, Trouche S, McNamara CG, Allen K & Dupret D. Hippocampal offline reactivation consolidates recently formed cell assemblies patterns during sharp-wave ripples. Neuron. 92(5),968–974 (2016).

21. Kudrimoti HS, Barnes CA & McNaughton BL. Reactivation of Hippocampal Cell Assemblies: Effects of Behavioral State, Experience, and EEG Dynamics. J Neurosc. 19(10):4090–101 (1999).

22. Girardeau G, Inema I & Buzsaki G. Reactivations of emotional memory in the hippocampus-amygdala system during sleep. Nat. Neurosci. 20, 1634–1642 (2017).

23. Lopes-dos-Santos V, Ribeiro S & Tort ABL. Detecting cell assemblies in large neuronal populations. J Neurosc Methods. 220(2), 149–166 (2013).

24. Stark E, Roux L, Eichler R & Buzsaki G. Local generation of multineuronal spike sequences in the hippocampal CA1 region. Proc Natl Acad Sci. 112(33), 10521–6 (2015).

25. Oliva A, Fernández-Ruiz A, Fermino de Oliveira E & Buzsáki G. Origin of gamma frequency power during hippocampal sharp-wave ripples. Cell Rep. 25(7), 1693–1700 (2018).

26. Sadowski JHLP, Jones MW & Mellor JR. Sharp-wave ripples orchestrate the induction of synaptic plasticity during reactivation of place cell firing patterns in the hippocampus. Cell Rep. 14(8): 1916–1929 (2016).

27. King C, Henze DA, Leinekugel X & Buzsaki G. Hebbian modification of a hippocampal population pattern in the rat. Journal of Phys. https://doi.org/10.1111/j.1469-7793.1999.00159.x (2004).

28. Grosmark AD & Buzsaki G. Diversity in neural firing dynamics supports both rigid and learned hippocampal sequences. Science. 351(6280),1440–1443 (2016).

29. Wagatsuma A, Okuyama T, Sun C, Smith LM, Abe K & Tonegawa S. Locus coeruleus input to hippocampal CA3 drives single-trial learning of a novel context. Proc Natl Acad Sci. 115 (2), 310–316 (2018).

30. Donegan ML, Stefanini F, Meira T, Gordon JA, Fusi S & Siegelbaum SA. Coding of social novelty in the hippocampal CA2 region and its disruption and rescue in a mouse model of schizophrenia. bioRxiv 833723; doi: https://doi.org/10.1101/833723

31. Chevaleyre V & Siegelbaum SA. Strong CA2 pyramidal neuron synapses define a powerful disynaptic cortico-hippocampal loop. Neuron 66 (4), 560–572 (2010).

32. Omer DB, Maimon SR, Las L, Ulanovsky N. Social place-cells in the bat hippocampus. Science. (359) 6372, 218–224 (2018).

33. Danjo T, Toyoizumi T, Fujisawa S. Spatial representations of self and other in the hippocampus. Science. (359) 6372, 213–218 (2018).

34. Rotenberg A and Muller RU. Variable place-cell coupling to a continuously viewed stimulus: evidence that the hippocampus acts as a perceptual system. Philosophical Transactions B. 352 (1360): 1505–1513 (1997).

35. Knierim JJ, Kudrimoti HS and McNaughton BL. Interactions between idiothetic cues and external landmarks in the control of place cells and head direction cells. Journal of Neurophysiology. 80 (1): 425–446 (1998).

36. Quiroga RQ, Reddy L, Kreiman G, Koch C & Fried I. Invariant visual representation by single neurons in the human brain. Nature. 435(7045), 1102–7 (2016).

37. Davoudi H & Foster DJ. Acute silencing of hippocampal CA3 reveals a dominant role in place field responses. Nat Neuroscience. 22, 337–342 (2019).

38. Tirko NN, Eyrin KW, Carcea I, Mitre M, Chao MV, Froemke RC & Tsien RW. Oxytocin transforms firing mode of CA2 hippocampal neurons. Neuron. 100(3), 593–608 (2019).

39. Pagani JH, Zhao M, Cui Z, Williams Avram SK, Caruana DA, Dudek SM & Young WS. Role of vasopressin 1b receptor in rodent aggressive behavior and synaptic plasticity in hippocampal area CA2. Mol Psychiatry. 20(4), 490–499 (2015).

40. Rao RP, von Heimendahl M, Bahr V & Brecht M. Neuronal responses to conspecifics in the ventral CA1. Cell Reports. 27, 3460–3472 (2019).

41. Okuyama T, Kitamura T, Roy DS, Itohara S & Tonegawa S. Ventral CA1 neurons store social memory. Science. 353, 6307, 1536–1541 (2016).

## METHODS REFERENCES

42. Fernández-Ruiz A, Oliva A, Nagy GA, Maurer AP, Berényi A & Buzsáki G. Entorhinal-CA3 dual-input control of spike timing in the hippocampus by theta-gamma coupling. Neuron. 93(5), 1213–1226 (2017).

43. Ylinen A, Bragin A, Nádasdy Z, Jandó G, Szabó I, Sik A & Buzsáki G. Sharp wave-associated high-frequency oscillation (200 Hz) in the intact hippocampus: Network and intracellular mechanisms. J. Neurosci. 15, 30–46 (1995).

44. Fernández-Ruiz A, Makarov VA, Benito N & Herreras O. Schaffer-specific local field potentials reflect discrete excitatory events at gamma frequency that may fire postsynaptic hippocampal CA1 units. J. Neurosci. 32(15),5165–76 (2012).

45. Fernández-Ruiz A, S. Muñoz S, Sancho M, Makarova J, Makarov VA & Herreras O. Cytoarchitectonic and dynamic origins of giant positive local field potentials in the dentate gyrus. J. Neurosci. 33(39),15518–32 (2013).

46. Roux L, Hu B, Eichler R, Stark E & Buzsaki G. Sharp wave ripples during learning stabilize hippocampal spatial map. Nat. Neurosci. 20, 845–853 (2017).

47. Mathis A, Mamidanna P, Cury KM, Abe T, Murthy VN, Mathis MW & Bethge M. DeepLabCut: markerless pose estimation of user-defined body parts with deep learning. Nat. Neurosci. 21, 1281–1289 (2018).

48. Pachitariu M, Steinmetz N, Kadir S, Carandini M, Harris KD. Fast and accurate spike sorting of high-channel count probes with KiloSort. 30th Conference on Neural Information Processing Systems (NIPS 2016), Barcelona, Spain (2016).

49. Mizuseki K, Sirota A, Pastalkova E & Buzsáki G. Theta oscillations provide temporal windows for local circuit computation in the entorhinal-hippocampal loop. Neuron. 64(2),267–80 (2009).

50. Skaggs WE, McNaughton BL, Wilson MA & Barnes CA. Theta Phase Precession in Hippocampal Neuronal Populations and the Compression of Temporal Sequences. Hippocampus. 6,149–172 (1996).

51. Oliva A, Fernandez-Ruiz A, Buzsaki G & Berenyi A. Spatial coding and physiological properties of hippocampal neurons in the Cornu Ammonis subregions. Hippocampus. 26(12), 1593–1607 (2016).

